# Brain endothelial cell tissue-nonspecific alkaline phosphatase (TNAP) activity promotes maintenance of barrier integrity *via* the ROCK pathway

**DOI:** 10.1101/2021.03.25.437097

**Authors:** Divine C. Nwafor, Allison L. Brichacek, Wei Wang, Nina Bidwai, Christa L. Lilly, José Luis Millán, Candice M. Brown

**Affiliations:** Department of Neuroscience, School of Medicine, West Virginia University Health Science Center, Morgantown, WV 26506, USA; Rockefeller Neuroscience Institute, West Virginia University; Department of Microbiology, Immunology, and Cell Biology, School of Medicine, West Virginia University Health Science Center, Morgantown, WV 26506, USA; Department of Biostatistics, School of Public Health, West Virginia University Health Science Center, Morgantown, WV 26506, USA; Sanford-Burnham Prebys Medical Discovery Institute, La Jolla, CA 92037, USA

**Keywords:** Tissue-nonspecific alkaline phosphatase, *Alpl*, Barrier integrity, ROCK, Brain microvasculature, Cytoskeleton

## Abstract

Blood-brain barrier (BBB) dysfunction is a key feature in many neuroinflammatory diseases. Yet, no therapies exist to effectively mitigate BBB dysfunction. A strategy to bridge this knowledge gap requires an examination of proteins localized to brain microvascular endothelial cells (BMECs) and evaluating their role in preserving barrier integrity. Tissue-nonspecific alkaline phosphatase (TNAP) is highly abundant in brain microvascular endothelial cells (BMECs); however, its function in BMECs remains unclear. We hypothesized that a loss or inhibition of TNAP activity on BMECs would impair barrier integrity through increased cytoskeletal remodeling driven by the Rho-associated protein kinase (ROCK) pathway. First, we examined barrier integrity in hCMEC/D3 cells treated with a TNAP inhibitor (TNAPi) and in primary BMECs (pBMECs) *via* the conditional deletion of TNAP in endothelial cells. Our results showed that both pharmacological inhibition and genetic conditional loss of TNAP significantly worsened endothelial barrier integrity compared to controls. Next, we examined the mechanisms through which TNAP activity exerts a protective phenotype on BMECs. Our results showed that hCMEC/D3 cells treated with TNAPi displayed remarkable phalloidin and vimentin cytoskeletal remodeling compared to control. We then examined the role of ROCK, a key player in cytoskeletal remodeling. Our results showed that TNAPi increased the expression of ROCK 1/2. Furthermore, inhibition of ROCK 1/2 with fasudil mitigated TNAPi-induced and VE-cKO barrier dysfunction. Collectively, our results support a novel mechanism through which loss of TNAP activity results in cerebrovascular dysfunction, and selective modulation of TNAP activity in BMECs may be a therapeutic strategy to improve BBB function.

## Introduction

The blood-brain barrier (BBB) is a dynamic vascular interface that separates the brain parenchyma from systemic circulation [1]. The brain’s microvasculature differs from the peripheral vasculature due to its unique composition of brain capillary endothelial cells linked tightly together by junctional proteins, surrounding pericytes, basal lamina, and astrocyte end-foot processes [2]. At the BBB, brain capillary endothelial cell-cell junctions are critical for maintaining the integrity of the BBB *via* paracellular transport size selective restriction of molecules, toxin, and cells [3]. Furthermore, the expression of specific proteins, enzymes, and transporters on luminal and abluminal surfaces of brain capillary endothelial cells help to regulate blood-to-brain trafficking of certain molecules [4]. One example of these important proteins is the endothelial sphingosine-1-receptor-1 (S1Pr_1_). Yanagida *et al.,* showed that conditional endothelial knockout of S1Pr1 led to a size selective BBB leakiness to fluorescent tracers less than 3 kDa. Furthermore, BBB leakiness in the S1Pr1 endothelial knockout mice was coupled to altered subcellular distribution of junctional proteins [5]. Thus, a cross-examination of specific proteins, enzymes, and transporters localized to brain capillary endothelial cells may provide therapeutic insight that may mitigate BBB dysfunction and the consequential long-term cognitive impairment seen across many neuroinflammatory conditions [2,6].

Alkaline phosphatases (APs) are found in numerous tissues, and are thought to play an important role in regulating inflammation [7]. There are four AP isozymes in humans (gene name in italics): placental alkaline phosphatase (PLAP; *ALPP*), germ cell alkaline phosphatase (GCAP; *ALPPL2*), intestinal alkaline phosphatase (IAP; *ALPI*), and tissue-nonspecific alkaline phosphatase (TNAP; *ALPL*; also known as *Akp2 or Alpl* in mouse) [8]. Of the four AP isozymes, only TNAP is expressed in the brain tissue of humans and rodents [9]. Moreover, biochemical and histological studies have shown that brain microvessels are abundantly rich in TNAP [10,11]. Several studies have demonstrated a role for APs in catalyzing the hydrolysis of nucleotides (ATP, ADP, and AMP) to free adenosine and inorganic phosphates [12]. Similarly, APs are able to dephosphorylate inflammatory molecules such as lipopolysaccharide (LPS), Damage-associated molecular patterns (DAMPs), and Pathogen-associated molecular patterns (PAMPs) [13–15]. Free adenosine and dephosphorylation of LPS (dLPS) are suggested to support the anti-inflammatory role for APs in disease [12,16–18]. However, the molecular mechanisms through which APs maintain homeostasis *via* its putative anti-inflammatory function remain unclear. Earlier investigations from our group revealed that brain microvascular TNAP activity is decreased as early as 24 h post-sepsis, and this decrease is sustained up to 7 days post-sepsis. Furthermore, the decrease in brain microvascular TNAP activity at 7-days post-sepsis was coupled to increased immunoglobulin G (IgG) permeability and sustained neuroinflammation. Moreover, treatment of septic animals with a specific TNAP inhibitor decreased brain endothelial junctional protein claudin-5 expression compared to vehicle treated septic and naïve mice [2,19]. These results suggest a putative role for microvascular TNAP activity in maintaining vascular barrier integrity at the endothelium. Therefore, in this study we elucidated the mechanistic role for TNAP at the brain endothelium and examined the molecular and cellular pathways targeted by TNAP.

Investigating the role of brain endothelial TNAP has been challenging for decades due to TNAP’s ubiquitous expression in other tissue types such as the liver, kidney, spleen, lung, bone, and diverse brain cell types [8,20]. Furthermore, mice with a global knockout of TNAP (*Alpl*^−/−^) die within days after birth from seizures and rickets characteristics of hypophosphatasia [21]. Newly generated pharmacological agents such as the TNAP inhibitor (TNAPi) probe and the pharmacological derivative 5-((5-chloro-2-methoxyphenyl) sulfonamido) nicotinamide (SBI-425) have advanced our knowledge on TNAP function in recent years [22,23]; however, the use of these drugs *in vivo* makes it difficult to delineate brain endothelial TNAP function as opposed to other cell types. The availability of the *Alpl* floxed mouse allowed for a targeted approach to delineate cell specific function by Cre-lox recombination [24]. We utilized both pharmacological, a TNAP inhibitor (TNAPi), and genetic, endothelial TNAP conditional knockouts, to demonstrate a mechanistic role for brain endothelial TNAP in maintaining barrier integrity *in vitro* and *ex vivo*. Taken together, our results suggest that TNAP plays a critical role in maintaining barrier integrity *via* Rho-associated protein kinase (ROCK) mediated by cytoskeletal reorganization.

## Methods

### Cell lines

The hCMEC/D3 human cerebral microvascular endothelial cell line (D3 cells) is an immortalized endothelial line that retains BBB characteristics *in vitro* [25]. The cell line was purchased from Cedarlane Labs (Burlington, NC)

### Animals

All experiments were conducted in accordance with the National Institutes of Health Guide for the Care and Use of Laboratory Animals and were approved by the Institutional Animal Care and Use Committee at West Virginia University. Male wild-type (WT; *C57BL6/J;* Bar Harbor, ME, Catalog # 000664) mice were bred in West Virginia University Health Sciences Center vivarium facilities and used for sepsis and stroke experiments at 3-5 months old. Floxed *Alpl* (*Alpl*^*fl/fl*^) mice on a C57BL/6J genetic background were crossed with B6.FVB-Tg(Cdh5-cre)7Mlia/J (VE-Cadherin Cre, Bar Harbor, ME, Catalog # 006137) mice obtained from Jackson Labs. Creation of *Alpl*^*fl/fl*^ mice is described in [24] and creation of VE-Cadherin Cre mice is described in [26]. VE-Cadherin Cre and *Alpl*^*fl/fl*^ mice were crossed to produce mice with a conditional deletion of *Alpl* in the endothelium (VE-cKO) and littermate control mice (*Alpl*^*fl/fl*^). For genotyping, DNA was extracted from ear snips using the Purelink Genomic DNA Mini Kit (Invitrogen, Carlsbad, CA, USA), and PCR products were amplified by using a Veriti 96-well Thermal Cycler (Applied Biosystems, ThermoFisher Scientific, Waltham, MA) under the following conditions: 94◻°C for 1◻min, [(94◻°C for 30◻sec, 60◻°C for 30◻sec, 72◻°C for 45◻sec) × 40 cycles], then 72◻°C for 1◻min. VE-Cadherin Cre specificity was determined by the presence of a 700 bp product using the following primers; ACRE_F: 5’- GAACCTGATGGACATGTTCAGGGA -3’, and ACRE_R: 5’- CAGAGTCATCCTTAGCGCCGTAAA -3’ [26]. Confirmation of floxed *Alpl* sites was determined by the presence of a 263 bp product using the following primer set; Alpl^flox^_F: 5’- GTTGCGATGTGTGAAGATGTCCTCG -3’, and Alpl^flox^_R: 5’- CTTGGGCTTGCTGTCGCCAGTAAC -3’. An additional strain was employed. Red fluorescent protein (RFP) B6.Cg-*Gt(ROSA)26Sor* (Ai9) mice under a genetic C57BL/6J background were obtained from Dr. Eric Tucker in the Department of Neuroscience at West Virginia University; generation of these mice is described in [27]. All mice were group housed in environmentally controlled conditions with a reverse light cycle (12:12 h light/dark cycle at 21 ± 1°C) and provided food and water *ad libitum.*

### Cecal Ligation and Puncture (CLP)

The cecal ligation and puncture (CLP) model of polymicrobial sepsis was employed as previously described [19,28]. Briefly, C57BL/6J mice were anesthetized by the inhalation of 1-2% isoflurane and abdominal access was obtained via a midline incision. The cecum was ligated below the ileocecal valve, punctured twice with a 22◻G needle through and through, and placed back into the abdominal cavity. The abdominal muscle and skin layer were closed with 6-0 and 5-0 sutures (Ethilon, Cornelia, GA) respectively. Sham-operated animals had their cecum isolated and then returned to the peritoneal cavity without being ligated or punctured. One mL of sterile 0.9% saline was administered subcutaneously (s.c.) for fluid resuscitation in all experimental groups. Mice used for all experiments were euthanized at seven days post-CLP.

### Transient Middle Cerebral Artery Occlusion (tMCAO)

tMCAO surgery was performed under isofluorane anesthesia as previously described [29]. Briefly, male C57BL/6J mice were subjected to 60 min tMCAO using silicon coated sutures (Cat. #702334, Doccol Corporation, MA) followed by reperfusion. Body temperatures were controlled at 37◻±◻0.5◻°C during occlusion. Occlusion and reperfusion were verified in each animal by a Laser Speckle Imager (Moor Instruments, England). Bupivacaine (2◻mg/kg, s.c.) was administered to relieve pain after surgery. Mice were followed for seven days post-stroke and euthanized.

### Tissue Collecting and Processing

Mice were deeply anesthetized with isoflurane and transcardially perfused (Masterflex 7524-10, Cole-Parmer, Vernon Hills, IL) as described previously [19]. Briefly, blood was removed with 0.9% saline followed by perfusion and fixation with 4% chilled paraformaldehyde (PFA, Fisher Scientific, Pittsburgh, PA). Perfused brains were post-fixed in 4% PFA overnight at 4°C. On the following day, brains were rinsed in 0.01 M phosphate buffered saline (PBS) and incubated sequentially in 15% and 30% sucrose in PBS for 24 h. Next, brains were co-embedded in 15% gelatin for sectioning. The gelatin block was processed sequentially through 4% PFA for 24 h, 15% sucrose for 48 h, and 30% sucrose for 48 h. The block was then trimmed and placed in a −80°C freezer for 1 h. Sectioning was performed in the coronal plane at 35 μm on a sliding microtome (HM 450, ThermoFisher Scientific).

### Primary Brain Microvascular Endothelial Cell (pBMEC) Culture

Brain microvascular endothelial cells (BMECs) were cultured from male and female mice as previously described [28]. Briefly, 6-8 week old *Alpl*^*fl/fl*^ (n = 5) and VE-cKO (n = 5) were perfused with 0.01 M phosphate buffered saline (PBS). Cortices were dissected, homogenized, digested in papain and DNase I (Worthington Biochemical Corp, Lakewood, NJ) at 37◻°C for 1◻h. The homogenate was then centrifuged (1360 × g) for 10◻minutes, followed by myelin removal. The cell pellet was resuspended in endothelial cell growth medium (ECGM: F12 medium with 10% fetal bovine serum (FBS), endothelial growth supplement, ascorbate (2.5◻μg/ml), L-glutamine (4◻mM), and heparin (10◻μg/ml), and plated into four collagen-coated wells (calf skin collagen, Sigma Aldrich, Milwaukee, WI) of a six-well plate. Cultures were treated with fresh ECGM medium the next day followed by treatment with puromycin hydrochloride (4 μg/ml) with EGCM◻+◻FBS for 2.5 days. Cultures reached confluency after 5–7 days and were used for barrier function assays.

### Brain Endothelial Cell Barrier Function Assays

Barrier function assays were carried out as previously described for D3 cells [19] and in pBMECs [28]. Briefly, D3 cells were seeded onto 3 independent collagen-coated 16-well E-Plate PET arrays (ACEA Biosciences, San Diego, CA) at a concentration of 20,000 cells/well and loaded onto an xCelligence RTCA DP system (ACEA Biosciences) enclosed in a cell culture incubator. Once D3 cells reached confluence ~ 24 h after seeding, triplicate wells in each array were treated with 200 μl of the following: 0.3% dimethyl sulfoxide (DMSO), tissue-nonspecific alkaline phosphatase inhibitor (TNAPi 100 μM; Millipore, Temecula, CA), or TNF-α and IFN-γ (10 ng/mL; Sigma-Aldrich). In a second set of experiments D3 cells in triplicate wells were treated with 200 μl of the following: 0.3% DMSO, TNAPi (100 μM), fasudil (10 μM; Sigma-Aldrich), or TNAPi (100 μM) and fasudil (10 μM). Barrier function assays were also performed in pBMECs cultures isolated from *Alpl*^*fl/fl*^ and VE-cKO mice. Barrier function of pBMEC cultures was recorded both with and without treatments. *Alpl*^*fl/fl*^ and VE-cKO endothelial cell cultures were treated with vehicle or 10 μM fasudil (i.e. *Alpl*^*fl/fl*^ untreated, *Alpl*^*fl/fl*^ fasudil, VE-cKO untreated, and VE-cKO fasudil). Cell impedance or normalized cell impedance was recorded and analyzed with RTCA Software 2.0 (ACEA Biosciences). Normalized cell impedance is calculated by dividing cell impedance at the normalized time (i.e. when cells are treated) by the original cell impedance. Untreated for all experiments refers to cells treated with vehicle (i.e. ECGM medium for pBMECs and endothelial cell growth basal medium-2 (EBM-2) for D3 cells).

### In-Cell Western (ICW) Assay

The ICW assay was performed using the Odyssey Imaging System (LI-COR Biosciences, Lincoln, NE) as previously described [30]. Briefly, D3 cell cultures were grown in 96-well plates until they reached confluency. Thereafter, D3 cells were treated with vehicle, 0.3% DMSO, or TNAPi (100 μM) for 24 h. The following day, cells were fixed with 4% PFA then permeabilized with 0.5% Triton X-100 for 15 min at room temperature and blocked with LI-COR Odyssey Blocking Solution (LI-COR Biosciences) for 1 h. The cells were then incubated overnight at 4°C with primary antibodies. The following primary antibodies were used: ROCK1 (Invitrogen (1:1000), AB_11155392), ROCK2 (Invitrogen (1:1000), AB_11157047), and RhoA (Abcam (1:1000), AB_10675086, Cambridge, MA). The next day, the cells were washed three times with PBS and incubated with the appropriate secondary IgG IRDye™ 800/680 antibody (1:10,000 dilution, LI-COR Biosciences) at room temperature for 2 h. The 96-well plates were scanned with the Odyssey CLx Infrared Imaging System (LI-COR Biosciences), and the integrated fluorescence intensities representing the protein expression levels were acquired using the Odyssey software (Odyssey Software Version 3.0, LI-COR Biosciences). The relative amount of the protein of interest was obtained by normalizing to total cell number (CellTag700 stain) in all experiments.

### Tissue-Nonspecific Alkaline Phosphatase (TNAP) Enzyme Activity Histochemistry

Brain tissue sections and cell cultures were evaluated for alkaline phosphatase enzyme activity with the BCIP/NBT substrate kit (SK-5400, Vector Laboratories, Burlingame, CA) as previously described [19,28]. Tissue sections and cells were rinsed three times in 0.1M Tris-HCl (pH = 9.5) for 5 min and incubated in BCIP/NBT staining solution for 4 h at room temperature. Following incubation, sections were rinsed in 0.01 M PBS and mounted onto microscope slides (Unifrost+, Azer Scientific, Morgantown, PA), air-dried overnight, dehydrated through a standard dehydration series, and cover-slipped with Permount (Fisher Scientific, Pittsburgh, PA).

### Immunohistochemistry

Brain sections and cell cultures were immunostained using standard immunohistochemistry techniques [19,28]. Briefly, tissue sections and cells were washed three times, permeabilized, and blocked for 30 min on a shaker. Tissue sections and cells were then incubated for 24 h with primary antibodies or 1 h in Alexa 594 phalloidin dye at room temperature, followed by a 2 h incubation with the appropriate secondary antibody at room temperature. The following primary antibodies were used with working dilutions and antibody identification indicated in parentheses: CD31 (RnD Systems (1:500), AB_1026192, Minneapolis, MN), Phalloidin (Invitrogen (1:1000), AB_2315633, Carlsbad, CA), and Vimentin (Cell Signaling Technologies (1:500), AB_10695459, Danvers, MA).

### Image Analysis

Sections were viewed on a Leica DM6B microscope (Leica Camera, Allendale, NJ) and images were captured using Leica LASX software (Leica Microsystems, Buffalo Grove, IL). Cell culture images were captured using the EVOS FL Auto 2.0 microscope (Thermofisher). The cortex, striatum, and hippocampus were identified by referring to the Allen Institute Brain Atlas (http://mouse.brain-map.org). TNAP and/or CD31 images were captured in the cortex, striatum, and hippocampus (40X magnification). 6 fields from 2 sections per animal were collected for the quantification of regional TNAP enzyme activity. CD31 and RFP images were captured in the cortex at 20X magnification. 30 images/cell culture well (n=3) were used for the quantification of phalloidin, vimentin, and TNAP enzyme activity stains. Images were quantified using FIJI/Image J version 2.0 software.

### Statistical Analysis

All experiments were executed to enhance rigor and avoid experimenter bias. Investigators were blinded to the experimental groups for all image analyses. Immunohistochemistry and ICW images were analyzed using a one-way analysis of variance (ANOVA) followed by Tukey’s multiple comparisons *post-hoc* test. The D3 cell barrier function assay was analyzed using a one-way repeated measure analysis of variance (ANOVA) followed by Tukey’s multiple comparisons *post-hoc* test. The pBMECs (i.e. *Alpl*^*fl/fl*^ versus VE-cKO) barrier function assay was analyzed using a two-way repeated measure analysis of variance (ANOVA). Whole brain AP activity assay in *Alpl*^*fl/fl*^ versus VE-cKO was analyzed using a two-tailed unpaired Student’s t-test. Barrier function assay for the fasudil treated pBMECs was analyzed using linear mixed modeling with repeated subject set to cell line and covariance matrix set to compound symmetry. The model effects included the time block (pre-treatment, fasudil treatment, and 48 h post-fasudil treatment), group (*Alpl*^*fl/fl*^ untreated, *Alpl*^*fl/fl*^ fasudil, VE-cKO untreated, and VE-cKO fasudil), and time block by group interaction. Comparisons within time blocks were made across groups on the least square means (LS means) with Tukey-Kramer adjustments. All analyses were conducted using SAS 9.4 (SAS software, Cary, NC) and GraphPad Prism 8.1 (GraphPad Software, La Jolla, CA). All *p*-values and *n* values are indicated in the figure legends. Results were expressed as means◻±◻SEM and *p*-values <0.05 were considered significant.

## Results

### Brain Microvascular TNAP Activity is Decreased in Models of Cerebrovascular Dysfunction

We have previously shown that brain microvascular TNAP activity is decreased at 24 h post-sepsis, and this decrease in TNAP activity is sustained up to 7 days post-sepsis [19,2]. Therefore, we examined whether the decrease in brain microvascular TNAP activity post-sepsis (Fig. 1a) extends to other neuroinflammatory conditions such as stroke. Cortex and striatum were examined because these regions revealed increased microvascular TNAP activity compared to other regions like the hippocampus in mice (Supplementary Fig. 1). Our results showed that following transient middle cerebral artery occlusion, the penumbra in the ipsilateral cortex and striatum exhibited decreased TNAP activity compared to the contralateral cortex and striatum (Fig. 1b). These results are consistent with an earlier study from our group showing that the loss of brain microvascular TNAP activity post-sepsis did not result from the loss of brain microvessels (Supplementary Fig. 2a) [19]. Similarly, CD31 and TNAP activity double-label histology in the penumbra of stroke tissue (striatum shown) revealed a loss of brain microvascular TNAP activity that is independent of CD31 positive vessel loss (Supplementary Fig. 2b). The loss of TNAP activity on CD31 positive microvessels after stroke and sepsis led us to speculate that TNAP may play an important role in cerebrovascular inflammation.

**Fig. 1.**
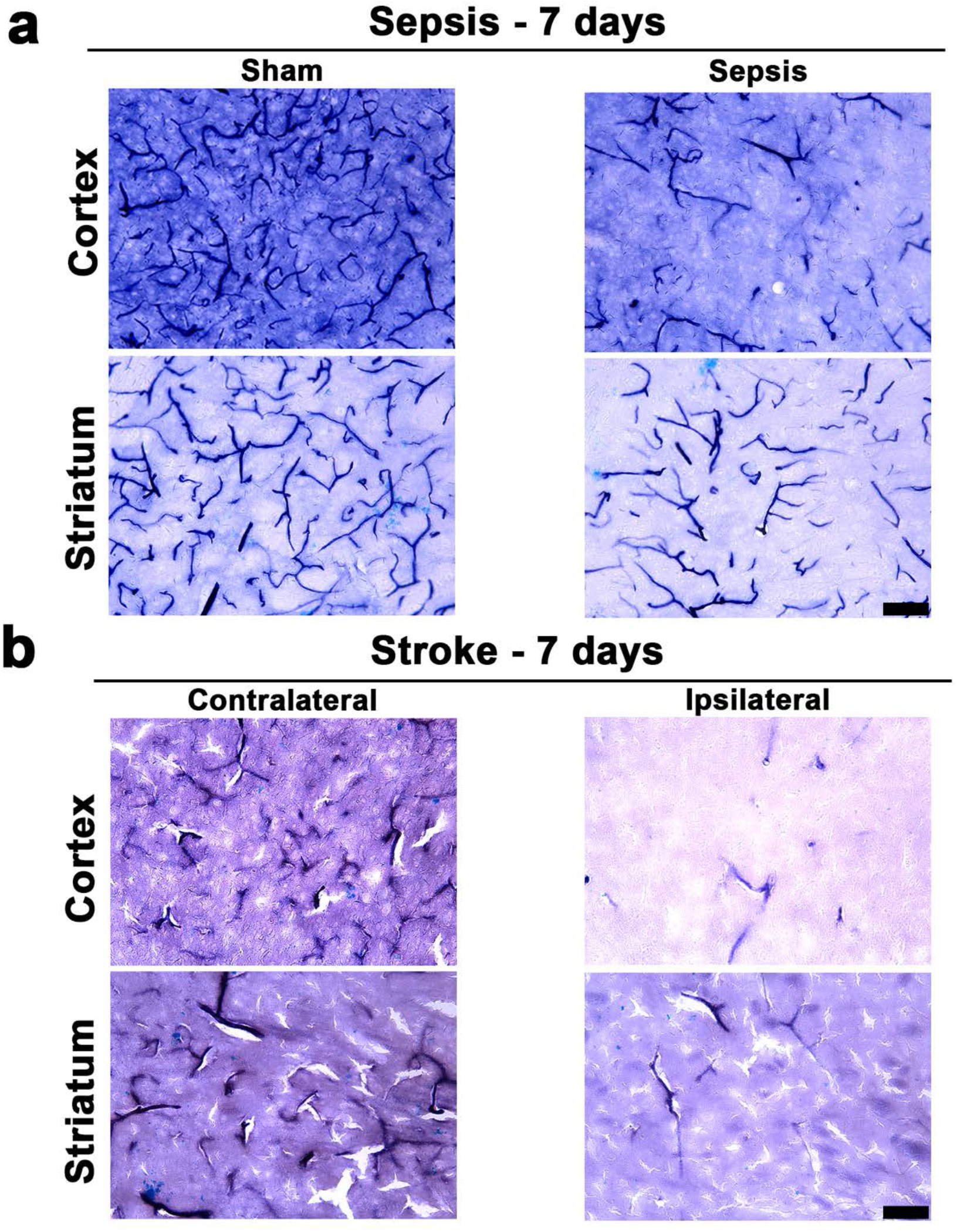
TNAP activity is decreased in sepsis and stroke (AD). (**a-b**) Representative images showed that TNAP activity in the cortex and striatum is decreased in sepsis (7 days post-sepsis) and stroke (7 days post-stroke) compared to appropriate controls. Images taken at 20X magnification and scale bar = 75 μm

### TNAP Inhibition in Brain Microvascular hCMEC/D3 Endothelial Cells Promotes Loss of Barrier Integrity

Brain microvascular endothelial cells are a key component of cerebral microvessels and play an important role in maintaining barrier integrity [31]. Next, we investigated whether a decrease in TNAP activity in brain endothelial cells promotes barrier dysfunction. Brain microvascular D3 endothelial cells were treated with TNAPi alone (a highly specific *in vitro* TNAP inhibitor [32]), TNF-α and IFN-γ, or TNF-α and IFN-γ combined with TNAPi. Our results showed that treatment of D3 endothelial cells with TNAPi (*p* = 0.01) and TNF-α and IFN-γ (*p* = 0.0005) significantly decreased endothelial TNAP activity compared to appropriate controls (Tukey’s multiple comparisons test, one-way ANOVA). Furthermore, a combined treatment of TNAPi and TNF-α and IFN-γ significantly decreased (*p* < 0.0001) endothelial TNAP activity compared to TNAPi alone (Tukey’s multiple comparisons test, one-way ANOVA) (Fig. 2a, b). We then examined whether the loss of TNAP activity in D3 endothelial cells following treatment with TNAPi or TNF-α and IFN-γ resulted in a loss of barrier function. Our results showed that both TNAPi (*p* < 0.0001) and TNF-α and IFN-γ (*p* < 0.0001) significantly decreased barrier integrity (impedance) compared to control (Tukey’s multiple comparisons test, one-way ANOVA) (Fig. 2c).

### Inhibition of Brain Endothelial TNAP Activity Induces Cytoskeletal Remodeling

The next set of studies sought to explore the mechanisms responsible for TNAPi-induced barrier dysfunction shown in **Figure 2**. Earlier studies by Deracinois *et al.* with a pan-phosphatase inhibitor (levamisole) suggested that the TNAP-dependent loss of barrier integrity in bovine capillary endothelial cells was associated with cytoskeleton remodeling [33]. However, levamisole is a non-specific inhibitor of TNAP and is capable of decreasing the activity of other phosphatases [34,23]. Hence, we examined cytoskeletal remodeling by using TNAPi. To do this, we treated D3 endothelial cells with TNAPi alone, TNF-α and IFN-γ, or TNF-α and IFN-γ combined with TNAPi. Thereafter, we immunostained for F-actin using phalloidin and intermediate filaments using vimentin 24 h following treatment. Our results showed that treatment of D3 endothelial cells with TNAPi alone (*p* = 0.0008) and TNF-α and IFN-γ (*p* < 0.002) significantly decreased F-actin fluorescence intensity compared to DMSO control (Tukey’s multiple comparisons test, one-way ANOVA). Furthermore, combined treatment of TNF-α and IFN-γ with TNAPi significantly decreased F-actin fluorescence intensity compared to TNAPi alone (*p* < 0.006) or TNF-α and IFN-γ (*p* = 0.0001; Tukey’s multiple comparisons test, one-way ANOVA). Moreover, we observed increased cell detachment (white arrows) in the TNAPi alone, TNF-α and IFN-γ, and TNF-α and IFN-γ combined with TNAPi groups compared to controls (i.e. DMSO or untreated) (Fig. 3a, b). Likewise, intermediate filament (i.e. vimentin) fluorescence intensity was significantly decreased in the TNAPi alone (*p* = 0.03) and TNF-α and IFN-γ combined with TNAPi (*p* = 0.008) when compared to DMSO control (Tukey’s multiple comparisons test, one-way ANOVA). However, treatment with TNF-α and IFN-γ alone did not appear to significantly alter the fluorescence intensity of vimentin (Fig. 3c, d).

**Fig. 2.**
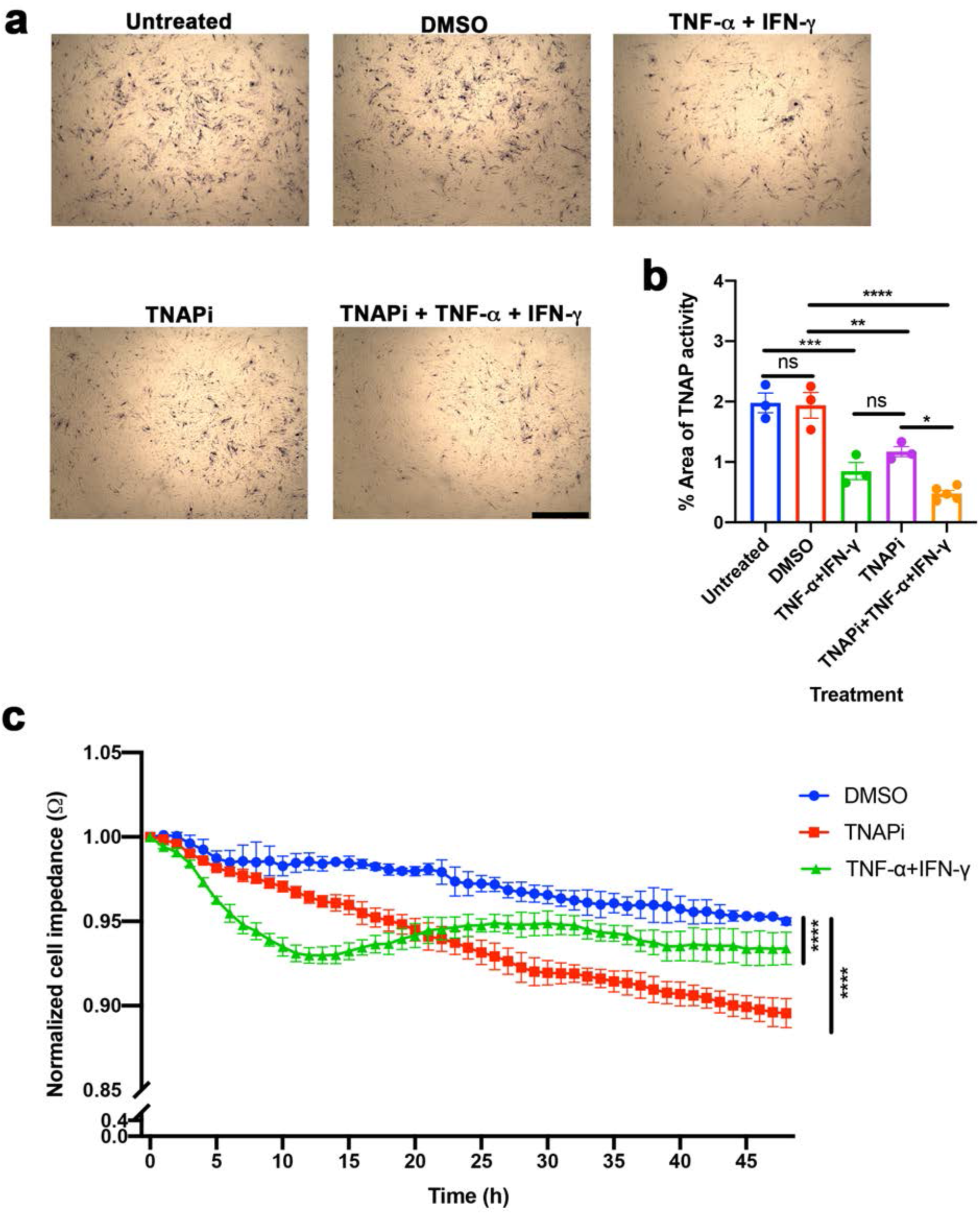
Inhibition of brain hCMEC/D3 endothelial TNAP promotes barrier dysfunction. (**a, b**) TNAPi (*p* = 0.01, Tukey’s multiple comparisons test, one-way ANOVA) and TNF-α and IFN-γ (*p* = 0.0005, Tukey’s multiple comparisons test, one-way ANOVA) significantly decreased brain endothelial TNAP activity compared to appropriate controls (DMSO or untreated). Moreover, combined treatment of TNAPi, TNF-α and IFN-γ significantly decreased (*p* < 0.0001, Tukey’s multiple comparisons test, one-way ANOVA) brain endothelial TNAP activity compared to TNAPi alone. (**c**) TNAPi (*p* < 0.0001, Tukey’s multiple comparisons test, repeated one-way ANOVA) and TNF-α and IFN-γ (*p* < 0.0001, Tukey’s multiple comparisons test, repeated one-way ANOVA) significantly decreased barrier integrity (impedance) compared to DMSO control. * indicates *p* < 0.05, \*\**p* < 0.01 \*\*\**p* < 0.001, and \*\*\*\**p* < 0.0001, and is considered significant. All data presented as mean ± SEM. Images taken at 10X magnification and scale bar = 1000 μm. n = 3-5 wells/treatment group, ns = not significant

**Fig. 3.**
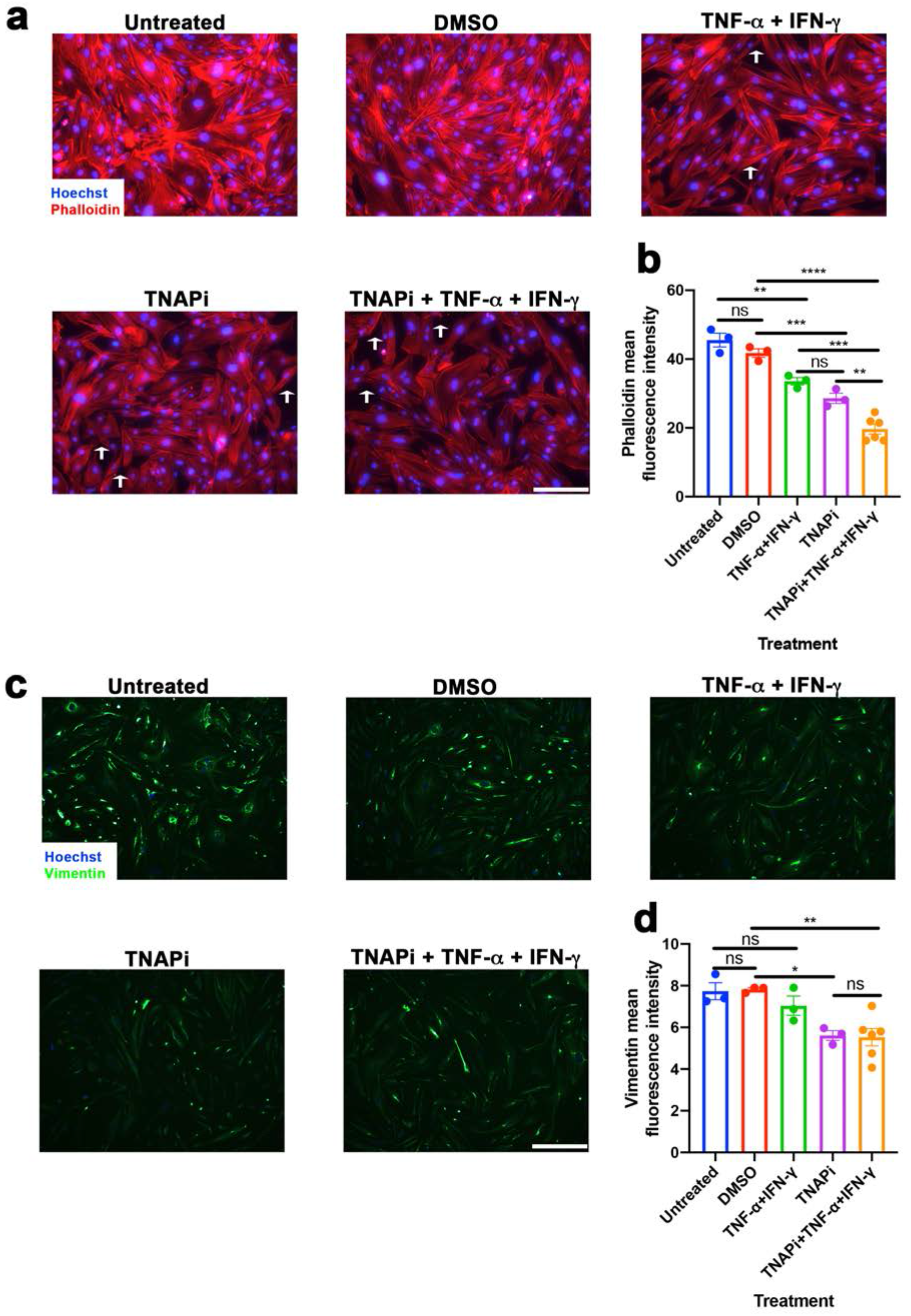
TNAPi induces cytoskeletal remodeling in hCMEC/D3 brain endothelial cells. (**a, b**) TNAPi (*p* = 0.0008, Tukey’s multiple comparisons test, one-way ANOVA) and TNF-α and IFN-γ (*p* < 0.002, Tukey’s multiple comparisons test, one-way ANOVA) significantly decreased phalloidin (F-actin) fluorescence intensity compared to appropriate controls (DMSO or untreated). Furthermore, combined treatment of TNAPi, TNF-α and IFN-γ significantly exacerbated the decrease in phalloidin fluorescence intensity compared to TNAPi (*p* < 0.006, Tukey’s multiple comparisons test, one-way ANOVA) or TNF-α and IFN-γ alone (*p* = 0.0001, Tukey’s multiple comparisons test, one-way ANOVA). Increased endothelial cell detachment (gaps, white arrows) accompanied the decrease in phalloidin fluorescence intensity. (**c, d**) TNAPi (*p* = 0.03, Tukey’s multiple comparisons test, one-way ANOVA) and combined treatment of TNAPi, TNF-α and IFN-γ (*p* = 0.008, Tukey’s multiple comparisons test, one-way ANOVA) significantly decreased vimentin fluorescence intensity compared to DMSO control. Interestingly, TNF-α and IFN-γ alone did not alter vimentin fluorescence intensity. * indicates *p* < 0.05, \*\**p* < 0.01 \*\*\**p* < 0.001, and \*\*\*\**p* < 0.0001, and is considered significant. All data presented as mean ± SEM. Images taken at 20X magnification and scale bar = 200 μm. n = 3-6 wells/treatment group, ns = not significant

### ROCK Protein Expression is Increased Following TNAP Inhibition in Endothelial Cells

Numerous studies have implicated the Rho/ROCK pathway as a critical mediator involved in endothelial cytoskeletal remodeling [35]. Based on the results derived in **Figure 3**, we employed in-cell Westerns (ICW) to examine the implications of TNAP inhibition on RhoA, ROCK 1, and ROCK 2 endothelial protein expression. Although RhoA expression increased relative to the appropriate control (DMSO) following treatment with TNAPi; however, this increase was not significant (Fig. 4a). Next, we examined downstream proteins (i.e. ROCK 1 and ROCK 2) involved in the Rho/ROCK pathway. Our results showed that ROCK 2 (*p* = 0.0003) and ROCK 1 (*p* = 0.02) protein expression was significantly increased following TNAPi treatment compared to DMSO control (Tukey’s multiple comparisons test, one-way ANOVA) (Fig. 4c-f). Interestingly, DMSO, the vehicle for TNAPi, significantly increased (*p* = 0.02) ROCK1 protein expression relative to untreated endothelial cells (Tukey’s multiple comparisons test, one-way ANOVA). This finding was not uncommon since DMSO has been shown to induce slight inflammation [36].

**Fig. 4.**
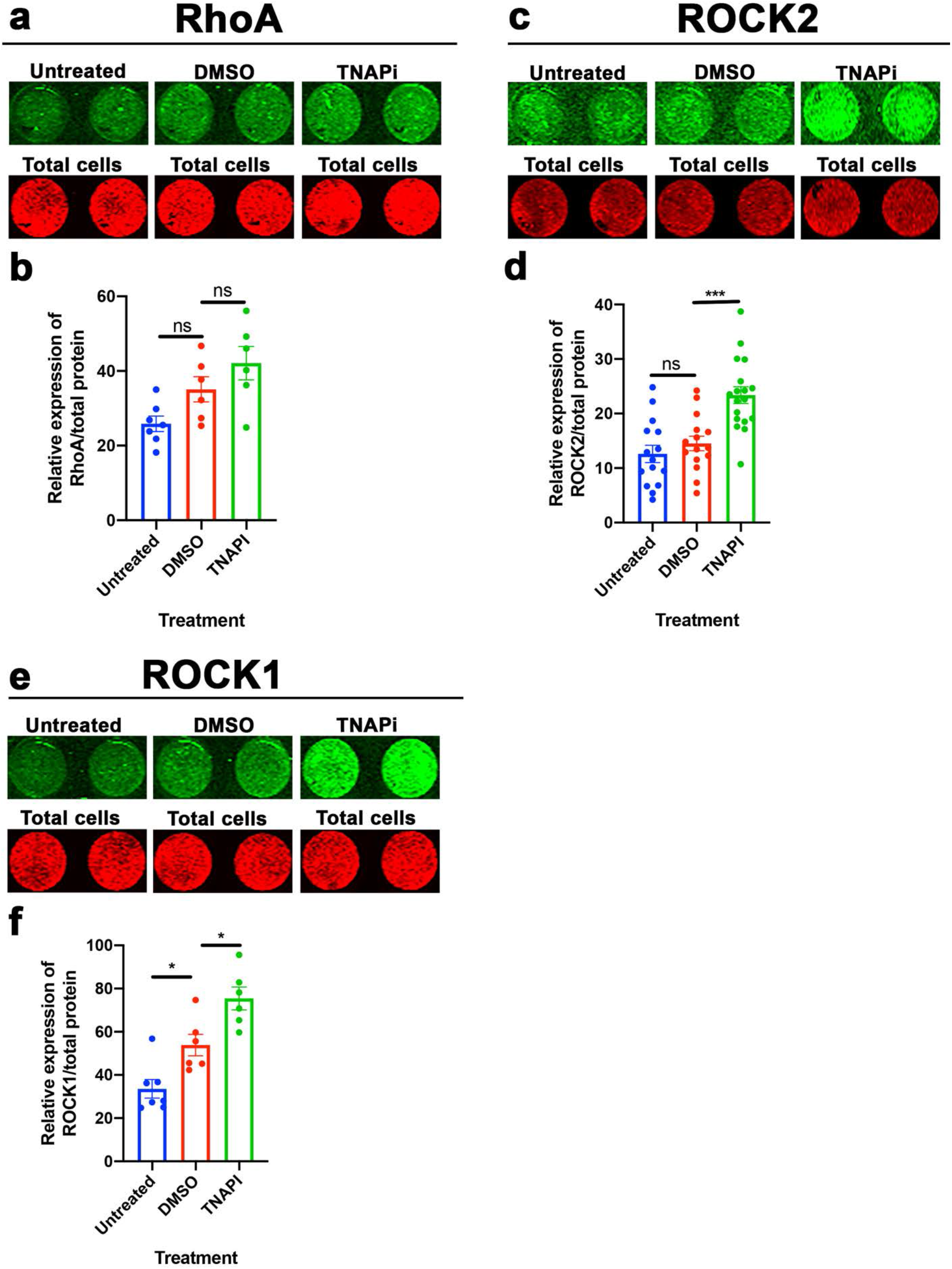
ROCK isoforms are increased following TNAPi treatment in hCMEC/D3 brain endothelial cells. (**a, b**) Rho protein was increased following treatment with TNAPi; however, this increase was not significant (*p* = 0.34, Tukey’s multiple comparisons test, one-way ANOVA). (**c-f**) Treatment with TNAPi increased ROCK 2 (*p* = 0.0003, Tukey’s multiple comparisons test, one-way ANOVA) and ROCK 1 (*p* = 0.02, Tukey’s multiple comparisons test, one-way ANOVA) protein expression compared to DMSO control. Fluorescence signal was normalized to total cell number for each respective well. * indicates *p* < 0.05 and \*\*\**p* < 0.001, and is considered significant. All data presented as mean ± SEM. Average n = 10 wells/treatment group, ns = not significant

### A ROCK Inhibitor Mitigates Loss of Barrier Integrity Following TNAP Inhibition

Fasudil is a potent competitive inhibitor of ROCK 1 and ROCK 2 [37]. Therefore, we examined whether fasudil alleviates TNAPi-induced barrier dysfunction. To do this, D3 endothelial cells were treated with TNAPi, fasudil, and TNAPi and fasudil. Following treatment, barrier integrity was measured over a 48-h period. Our results showed that TNAPi treatment significantly decreased (*p* < 0.0001, Tukey’s multiple comparisons test, repeated one-way ANOVA) barrier integrity compared to DMSO control. A combination treatment of fasudil and TNAPi revealed a significantly improved (*p* < 0.0001, Tukey’s multiple comparisons test, repeated one-way ANOVA) barrier integrity relative to TNAPi alone (Fig. 5). These results implicate ROCK proteins as downstream mediators of TNAP inhibition.

**Fig. 5.**
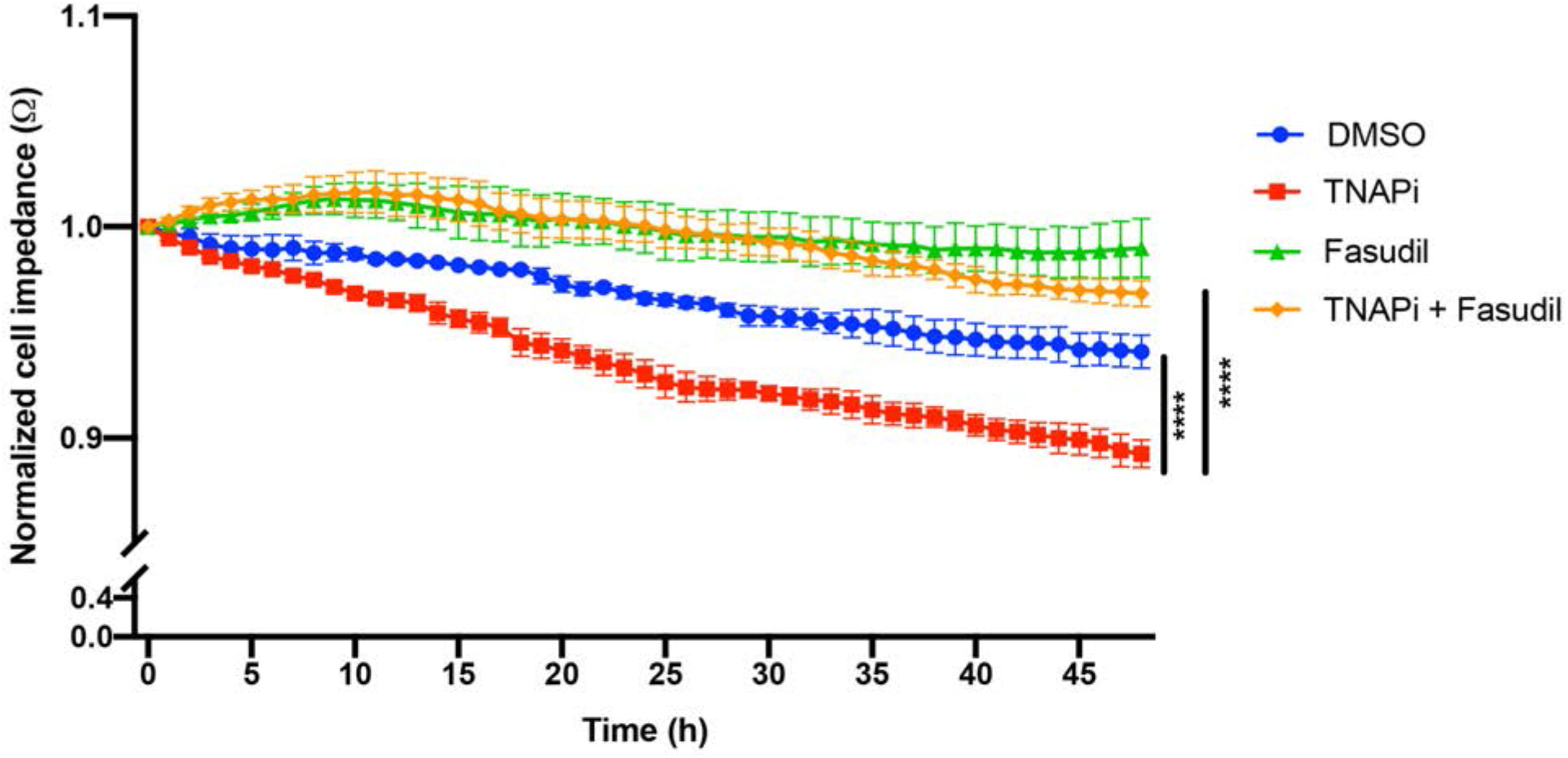
Fasudil mitigates TNAPi-induced barrier dysfunction in hCMEC/D3 brain endothelial cells. TNAPi alone significantly decreased (*p* < 0.0001, Tukey’s multiple comparisons test, repeated one-way ANOVA) barrier integrity compared to DMSO control. Combined treatment of TNAPi and fasudil significantly (*p* < 0.0001, Tukey’s multiple comparisons test, repeated one-way ANOVA) rescues TNAPi-induced barrier dysfunction. **** indicates *p* < 0.001, and is considered significant. All data presented as mean ± SEM. n = 3 wells/treatment group

### Conditional Deletion of TNAP in Primary Brain Microvascular Endothelial Cells Reduces Barrier Integrity

To further establish an important role for endothelial TNAP in barrier maintenance, we generated a mouse model with a conditional deletion of endothelial *Alpl* - the gene that encodes TNAP – in endothelial cells [38]. To do this, we bred a *Cdh5*-Cre driver mouse with the previously published *Alpl* floxed mouse [24]. *Cdh5* encodes VE-cadherin, a marker of endothelial cells. First, we verified that Cre expression was specific for vascular endothelial cells by crossing the *Cdh5*-Cre mice with Ai9 RFP promoter mice (Supplementary Fig. 4). Then, we measured whole brain AP activity in offspring littermates. VE-cKO (endothelial TNAP knockouts) mice revealed a significantly decreased (*t* = 5.4, *p* = 0.006, unpaired t-test) whole brain AP activity compared to *Alpl*^*fl/fl*^ mice (littermate control) (Fig. 6a). To further demonstrate the loss of TNAP in endothelial cells, we isolated and cultured primary brain microvascular endothelial cells (pBMECs) from *Alpl*^*fl/fl*^ and VE-cKO mice. Our results revealed an absence of TNAP activity in VE-cKO pBMECs compared to *Alpl*^*fl/fl*^ pBMECs (Fig. 6b). Consequently, we assessed whether the loss of TNAP on VE-cKO pBMECs were comparable to the barrier assay results derived with TNAPi in D3 endothelial cells. Our results revealed a significant loss of barrier integrity (*p* < 0.0001) in VE-cKO pBMECs compared to *Alpl*^*fl/fl*^ pBMECs (Tukey’s multiple comparisons test, repeated two-way ANOVA) (Fig. 6c).

**Fig. 6.**
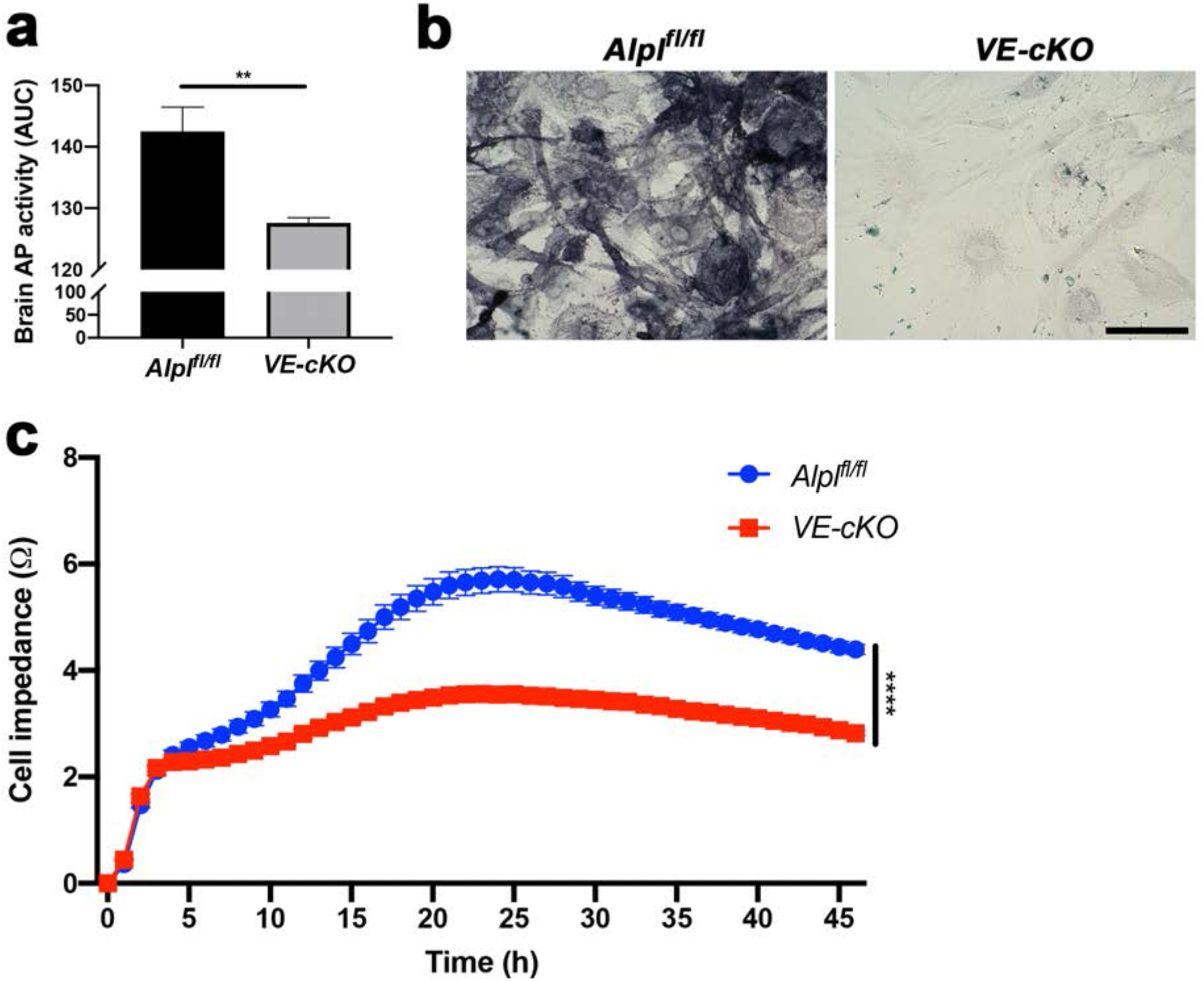
Conditional loss of endothelial TNAP worsens barrier integrity. (**a**) Whole brain AP activity is significantly increased (*p* = 0.006, unpaired t-test) in *Alpl*^fl/fl^ (n = 2, littermate control) mice compared to VE-cKO (n = 4, endothelial TNAP knockout) pBMECs. (**b**) VE-cKO pBMECs were absent of TNAP activity stain (purple) compared to *Alpl*^fl/fl^ pBMECs. (**c**) Barrier integrity in VE-cKO pBMECs was significantly decreased (*p* < 0.0001, Tukey’s multiple comparisons test, repeated two-way ANOVA) compared to *Alpl*^fl/fl^ pBMECs. ** indicates *p* < 0.01 and \*\*\*\**p* < 0.0001, and is considered significant. All data presented as mean ± SEM. Images taken at 40X magnification and scale bar = 100 μm. n = 4 wells/treatment group

### Fasudil Rescues Barrier Integrity in VE-cKO Endothelial Cells

Next, we assessed whether ROCK inhibition would mitigate the loss of barrier integrity shown in **Figure 6c**. Barrier integrity data was collected at three timepoints: pre-treatment, fasudil treatment, and 48 h post-fasudil treatment (i.e. drug elimination). Our results shown in **Figure 7a** (pre-treatment) paralleled the results derived in **Figure 6c** as expected (i.e. VE-cKO pBMECs demonstrated a significant loss of barrier integrity (*p* = 0.0009) compared to *Alpl*^*fl/fl*^ pBMECs (repeated three-way ANOVA). Following treatment with fasudil, barrier integrity in VE-cKO pBMECs significantly improved (*p* < 0.0001) compared to untreated VE-cKO pBMECs (Tukey-Kramer comparisons test, linear mixed modeling). Interestingly, the improvement of barrier integrity in fasudil treated VE-cKO pBMECs was comparable (*p* = 0.89) to untreated *Alpl*^*fl/fl*^ pBMECs (Tukey-Kramer comparisons test, linear mixed modeling) (Fig. 7b). 48 h post-fasudil treatment, barrier integrity in fasudil treated VE-cKO pBMECs declined and became comparable (*p* = 0.52) to untreated VE-cKO pBMECs (Tukey-Kramer comparisons test, linear mixed modeling) (Fig. 7c).

**Fig. 7.**
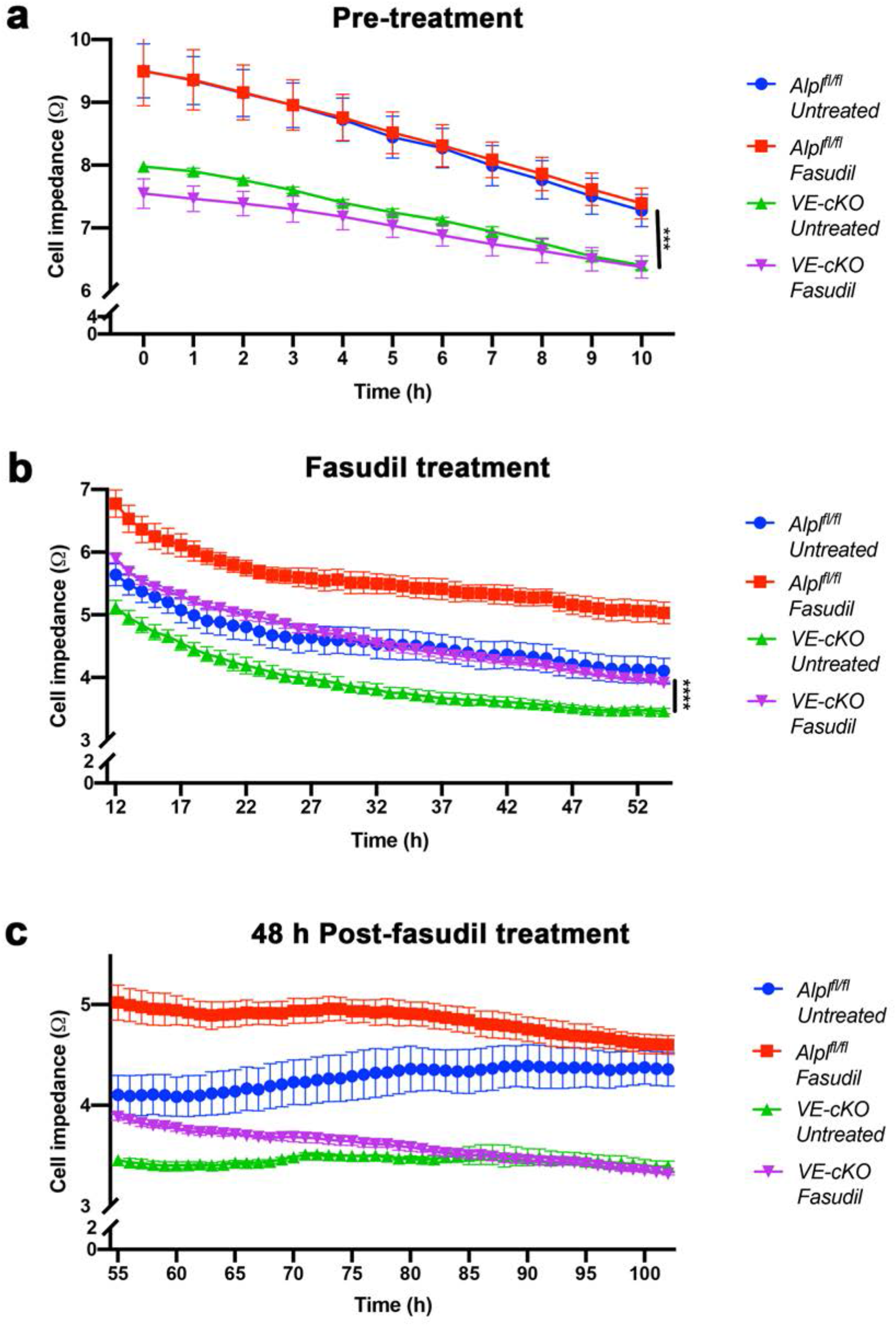
Fasudil mitigates barrier integrity loss in VE-cKO pBMECs. (**a**) Barrier integrity was significantly decreased (*p* = 0.0009, repeated three-way ANOVA) in VE-cKO pBMECs compared to *Alpl*^*fl/fl*^ pBMECs before fasudil treatment. (**b**) Fasudil treatment significantly mitigated (*p* < 0.0001, Tukey-Kramer comparisons test, linear mixed modeling) loss of barrier integrity in fasudil treated VE-cKO pBMECs compared untreated VE-cKO pBMECs. (**c**) 48 h post-fasudil treatment (i.e. drug elimination) barrier integrity in fasudil treated VE-cKO pBMECs was comparable (*p* = 0.52, Tukey-Kramer comparisons test, linear mixed modeling) to untreated VE-cKO pBMECs. *** indicates *p* < 0.001, and \*\*\*\**p* < 0.0001, and is considered significant. All data presented as mean ± SEM. n = 3 wells/treatment group

## Discussion

In the present study, we sought to elucidate the mechanistic role of an AP isozyme, TNAP, in brain microvascular endothelial cells. We have shown previously that microvascular TNAP activity is decreased as early as 24 h post-sepsis and sustained up to 7 days post-sepsis in the brains of mice [2,19]. In addition, we showed that the decrease in TNAP activity in brain microvessels extrapolates to other acute inflammatory conditions such as stroke. The observation of a loss microvascular TNAP activity in late (7 days) sepsis and ischemic stroke highlights a putative role for TNAP at the brain’s vasculature. Our results demonstrate an important mechanistic role for endothelial TNAP in maintaining barrier integrity by preventing the ROCK-mediated disruption of the cytoskeleton. Of note, it remains unclear how TNAP’s action on ROCK proteins converge into the well-known function of TNAP involving the dephosphorylation of nucleotides, LPS, DAMPs, and PAMPs [13–15]. Inflammatory molecules such as LPS have been shown in other models to increase ROCK protein expression and cause endothelial barrier integrity loss [39–41]; hence, it is likely that the action of LPS, DAMPs, and PAMPs are upstream relative to the novel mechanistic findings shown in this study. Future studies will address the convergence of these pathways.

Circulating levels of pro-inflammatory cytokines such as TNF-α and IFN-γ have been shown to be increased in neuroinflammatory conditions like sepsis and stroke [42–45]. Our laboratory has previously shown that there is a sustained increase in pro-inflammatory cytokines up to 7 days post-sepsis, and this increase is coupled to a loss of TNAP activity in brain microvessels [19]; therefore, we examined whether treatment with TNF-α and IFN-γ decreased endothelial TNAP activity similar to TNAPi. Our results showed that treatment of the brain microvascular D3 endothelial cells with TNF-α and IFN-γ sufficiently decreased TNAP activity similarly to TNAPi treatment alone. This finding demonstrates that pro-inflammatory cytokines may, in part, be responsible for the initial loss of brain microvascular TNAP activity seen in sepsis and stroke. Furthermore, we showed that the decrease in endothelial TNAP activity or the conditional loss of TNAP worsened endothelial cell barrier integrity. Owing to these findings, we propose that impaired BBB function seen in sepsis and stroke pathogenesis originates from endothelial cell damage, and the loss of endothelial TNAP activity contributes to BBB dysfunction.

The overall findings from our pharmacological and genetic studies have identified a novel molecular mechanism through which brain endothelial cell TNAP regulates BBB integrity. Importantly, our results substantially extend the findings from a previous study which showed that treatment of bovine capillary endothelial cells with a pan-AP inhibitor, i.e. levamisole, induced the retraction of endothelial cells [33]. Endothelial cell retraction and detachment are prototypical indicators of cytoskeletal reorganization and have been shown to accompany BBB dysfunction and increase cellular permeability [46,47]. We observed that treatment of BMECs with TNF-α and IFN-γ combined with inhibition of TNAP enzymatic activity disrupted the actin cytoskeleton. More importantly, the reorganization of the BMEC actin cytoskeleton was accompanied by increased cell detachment between adjacent endothelial cells, which represents reduced cell-cell junctional protein contact and adhesion. Junctional proteins such as claudin-5 are connected to the actin cytoskeleton *via* scaffold proteins and have been shown to play an important role in maintaining paracellular barrier permeability [48]. Therefore, it becomes plausible that the observed TNAPi-induced loss of endothelial barrier integrity shown in this study originates from the inability of brain endothelial cells to form proper cell-cell tight junctions. Therefore, we propose that the observed TNAPi-induced loss of endothelial barrier integrity shown in this study originates from the inability of brain endothelial cells to form proper cell-cell adhesion. This is supported by a previous *in vivo* study from our laboratory which showed that the junctional protein claudin-5 is decreased in the brains of septic mice treated intraperitoneally with an *in vivo* TNAP inhibitor, SBI-425, compared to vehicle treated septic mice [19]. In addition, morphological analyses of endothelial TNAP conditional knockout cell cultures with the AP activity stain revealed endothelial retraction and cellular detachment similar to TNAPi treated microvascular D3 endothelial cells (data not shown).

Intermediate filaments have been shown to directly and indirectly interact with actin [49,50]. This interaction is demonstrated by studies which showed that actin disruption affected intermediate filament sub-localization networks in cells [51–53]. We observed that the disruption of actin following TNAP enzyme inhibition disrupted the intermediate filament. Vimentin is one of the major intermediate filaments shown to provide a structural support for cells [54]. Interestingly, we only observed diminished vimentin fluorescence intensity following inhibition of TNAP activity, but not with TNF-α and IFN-γ treatment. In contrast, other studies that employed different cell types, human umbilical vein endothelial cells (HUVECs) or astrocytes, have demonstrated that TNF-α treatment increased vimentin protein expression [55,56]. We suggest that the properties of individual cell types, the dosage used, and use of TNF-α alone instead of TNF-α and IFN-γ are responsible for this discrepancy. Taken together, our results suggest that the sustained loss of microvascular TNAP activity induced by the presence of pro-inflammatory cytokines may be detrimental to the actin cytoskeleton and intermediate filaments. However, it remains unclear whether the loss of actin initiates vimentin disruption following TNAPi treatment or *vice versa*.

ROCK protein expression has been shown to play a role in regulating cytoskeletal proteins such as vimentin and actin by initiating F-actin contraction/retraction and cellular detachment [37,35,46]. Upstream of ROCK is the RhoA protein, one of many proteins that drives the activation of ROCK, and the Rho-ROCK pathway has been shown to play an important role in endothelial cell function Based on our cytoskeletal findings, we addressed whether the Rho/ROCK pathway was implicated in TNAP signaling. Our results showed that ROCK (1/2) protein expression was significantly increased following TNAPi treatment, along with a trending decrease in RhoA. protein levels. This finding indicates that TNAP-mediated signaling mechanism which suppresses ROCK activation is likely mediated by at least one separate pathway in addition to the canonical RhoA pathway. We speculate that the drastic effect of TNAP inhibition on ROCK 2 compared to ROCK 1 protein expression arises from the increased expression of ROCK 2 in brain tissue compared to ROCK 1 [37]. Furthermore, it is also likely that ROCK 1 and ROCK 2 may play independent effector roles downstream. For example, Shi *et el.*, showed that ROCK 1 regulates the actin cytoskeleton through myosin light chain 2 (MLC2) phosphorylation while ROCK 2 regulates the actin cytoskeleton through cofilin phosphorylation in mouse embryonic fibroblast (MEF) cells [57]. Finally, we utilized a ROCK1/2 inhibitor (fasudil) to further demonstrate the involvement of ROCK proteins in a novel TNAP signaling pathway in both D3 cells and pBMECs. These results contribute a novel mechanism in support of the neuroprotective and anti-inflammatory benefit of fasudil treatment established in preclinical models of sepsis and stroke [58–61].

BBB dysfunction is a common feature in many neuroinflammatory disorders [4]. Diminished BBB function is characterized by increased loss of junctional proteins, increased paracellular permeability of molecules and immune cells into the brain parenchyma, and endothelial cell transcytosis of immune cells into the parenchyma [2]. To our knowledge, this is the first study to describe the novel role played by endothelial TNAP in maintaining paracellular barrier integrity since our barrier assays can only assess this parameter. Our results suggest that endothelial TNAP exerts an inhibitory function on the activity of endothelial ROCK proteins in health and this function is dysregulated during acute injury as suggested by our septic and stroke studies. We propose a working model (Fig. 8) in which TNAP inhibition or loss prevents the contraction/retraction of endothelial cells and junctional protein disruption. During inflammation and/or acute injury, the rapid increase in pro-inflammatory cytokines leads to a reduction in endothelial TNAP activity on microvessels. The sustained loss of TNAP activity leads to cytoskeletal remodeling indicated by endothelial contraction and junctional protein detachment, which then permits the paracellular infiltration of pro-inflammatory molecules and immune cells that promote astrogliosis and microglial activation.

**Fig. 8.**
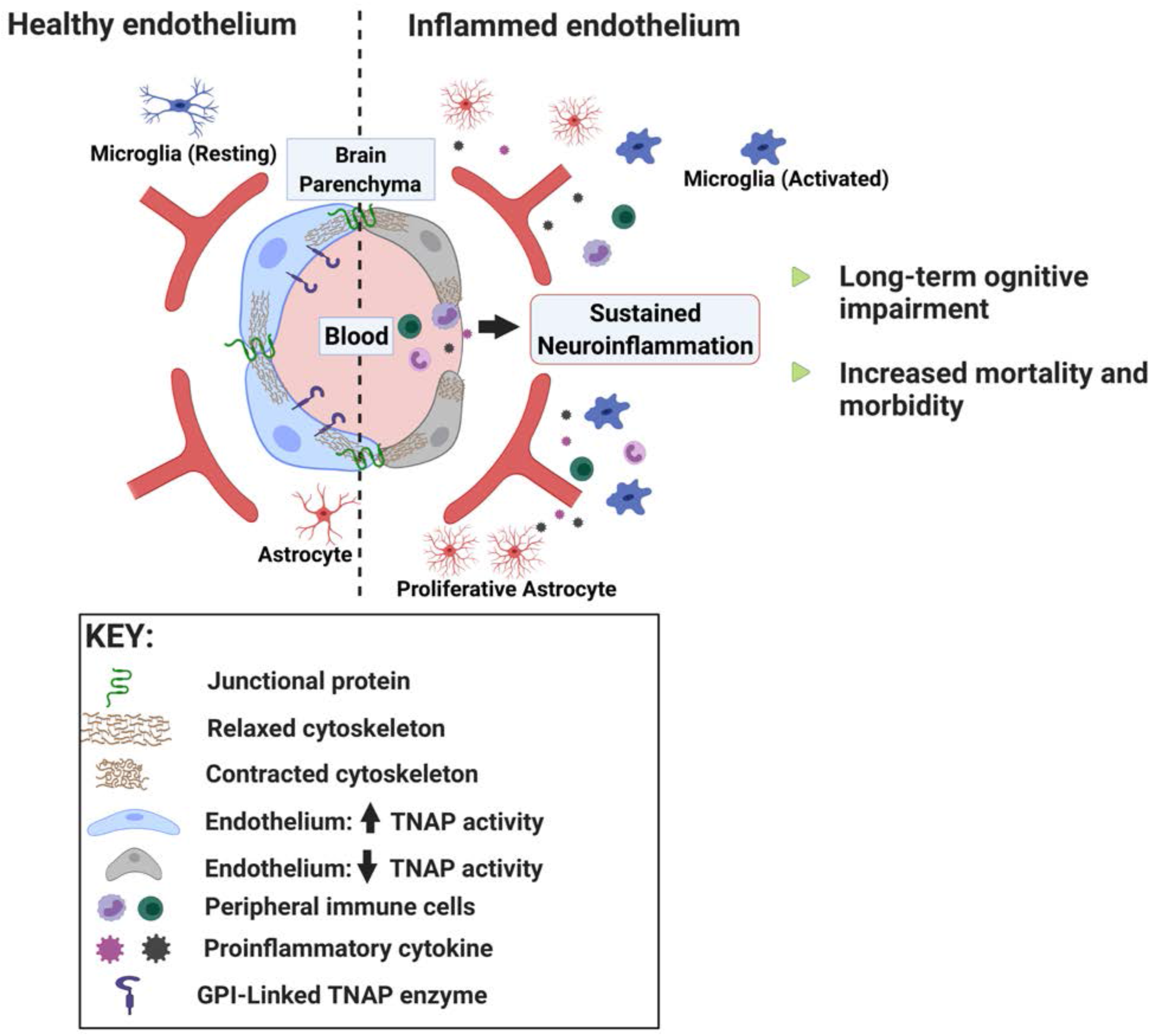
A proposed function of TNAP at the endothelium. In health, endothelial TNAP maintains barrier integrity *via* regulation of the actin cytoskeleton; however during inflammation, systemic pro-inflammatory cytokines decrease TNAP activity in endothelial cells. The decrease in endothelial TNAP activity promotes actin cytoskeleton contraction and detachment of adjacent endothelial cells which lead to a loss of barrier integrity. Thereafter, pro-inflammatory cytokines and immune cells infiltrate the brain parenchyma promoting the activation and proliferation of microglia and astrocytes. Sustained neuroinflammation ultimately leads to increased mortality and morbidity. Image credit: Biorender.com

Despite the current advances discussed in this study, some limitations exist. Since the experiments were carried out using *in vitro* and *ex vivo* methods, the contribution of astrocytes or pericytes is not clear and will be addressed in future studies. Since D3 cells were generated from a female donor and mixed sex pBMECs were used in the *ex vivo* experiments, we were unable to delineate sex-specific pathways. Future studies will address the role of sex on brain endothelial TNAP function. Furthermore, the assays used in this study were unable to delineate how endothelial TNAP may affect transcellular permeability. In summary, our results support a novel role for TNAP signaling in maintaining BMEC barrier integrity. Therapeutic strategies that modulate the endothelial TNAP activity may be beneficial for the treatment of BBB dysfunction or temporal - targeted drug delivery for various neurological disorders such as brain tumors, ischemic stroke, or Alzheimer’s disease.

## Acknowledgement

The authors gratefully thank the West Virginia University Office of Laboratory Animal Resources and the West Virginia University Experimental Stroke Core for their assistance with stroke surgeries.

## Author contributions

D.C.N, A.L.B, and C.M.B designed the studies. D.C.N. and W.W. performed all cell culture experiments. D.C.N. and N.B. performed image analyses. A.L.B. generated and genotyped *Alpl*^*fl/fl*^ and VE-cKO animals needed for experiments. D.C.N. and C.L.L. performed statistical analyses of data and generation of graphs. D.C.N. and C.M.B. wrote the manuscript. JL.M. provided the *Alpl*^*fl/fl*^ mice. All authors read and revised the final manuscript.

## Availability of data and material

Data available upon request from the corresponding author.

## Code availability

Not applicable

## References

1. Banks WA (2016) From blood-brain barrier to blood-brain interface: new opportunities for CNS drug delivery. Nat Rev Drug Discov 15 (4):275–292. doi:10.1038/nrd.2015.21

2. Nwafor DC, Brichacek AL, Mohammad AS, Griffith J, Lucke-Wold BP, Benkovic SA, Geldenhuys WJ, Lockman PR, Brown CM (2019) Targeting the Blood-Brain Barrier to Prevent Sepsis-Associated Cognitive Impairment. J Cent Nerv Syst Dis 11:1179573519840652. doi:10.1177/1179573519840652

3. Daneman R, Prat A (2015) The blood-brain barrier. Cold Spring Harb Perspect Biol 7 (1):a020412. doi:10.1101/cshperspect.a020412

4. Varatharaj A, Galea I (2017) The blood-brain barrier in systemic inflammation. Brain Behav Immun 60:1–12. doi:10.1016/j.bbi.2016.03.010

5. Yanagida K, Liu CH, Faraco G, Galvani S, Smith HK, Burg N, Anrather J, Sanchez T, Iadecola C, Hla T (2017) Size-selective opening of the blood-brain barrier by targeting endothelial sphingosine 1-phosphate receptor 1. Proc Natl Acad Sci U S A 114 (17):4531–4536. doi:10.1073/pnas.1618659114

6. Obermeier B, Daneman R, Ransohoff RM (2013) Development, maintenance and disruption of the blood-brain barrier. Nat Med 19 (12):1584–1596. doi:10.1038/nm.3407

7. Rader BA (2017) Alkaline Phosphatase, an Unconventional Immune Protein. Front Immunol 8:897. doi:10.3389/fimmu.2017.00897

8. Buchet R, Millan JL, Magne D (2013) Multisystemic functions of alkaline phosphatases. Methods Mol Biol 1053:27–51. doi:10.1007/978-1-62703-562-0_3

9. Brun-Heath I, Ermonval M, Chabrol E, Xiao J, Palkovits M, Lyck R, Miller F, Couraud PO, Mornet E, Fonta C (2011) Differential expression of the bone and the liver tissue non-specific alkaline phosphatase isoforms in brain tissues. Cell and tissue research 343 (3):521–536. doi:10.1007/s00441-010-1111-4

10. Farkas-Bargeton E, Arsenio-Nunes ML (1970) [Maturation of enzymatic equipment in the vessel walls of the nervous system. Histochemical study]. Acta Neuropathol 15 (3):251–271. doi:10.1007/BF00686771

11. Williams SK, Gillis JF, Matthews MA, Wagner RC, Bitensky MW (1980) Isolation and characterization of brain endothelial cells: morphology and enzyme activity. J Neurochem 35 (2):374–381. doi:10.1111/j.1471-4159.1980.tb06274.x

12. Picher M, Burch LH, Hirsh AJ, Spychala J, Boucher RC (2003) Ecto 5’-nucleotidase and nonspecific alkaline phosphatase. Two AMP-hydrolyzing ectoenzymes with distinct roles in human airways. J Biol Chem 278 (15):13468–13479. doi:10.1074/jbc.M300569200

13. Peters E, Geraci S, Heemskerk S, Wilmer MJ, Bilos A, Kraenzlin B, Gretz N, Pickkers P, Masereeuw R (2015) Alkaline phosphatase protects against renal inflammation through dephosphorylation of lipopolysaccharide and adenosine triphosphate. Br J Pharmacol 172 (20):4932–4945. doi:10.1111/bph.13261

14. Bentala H, Verweij WR, Huizinga-Van der Vlag A, van Loenen-Weemaes AM, Meijer DK, Poelstra K (2002) Removal of phosphate from lipid A as a strategy to detoxify lipopolysaccharide. Shock 18 (6):561–566. doi:10.1097/00024382-200212000-00013

15. Bates JM, Akerlund J, Mittge E, Guillemin K (2007) Intestinal alkaline phosphatase detoxifies lipopolysaccharide and prevents inflammation in zebrafish in response to the gut microbiota. Cell Host Microbe 2 (6):371–382. doi:10.1016/j.chom.2007.10.010

16. Peters E, Heemskerk S, Masereeuw R, Pickkers P (2014) Alkaline phosphatase: a possible treatment for sepsis-associated acute kidney injury in critically ill patients. Am J Kidney Dis 63 (6):1038–1048. doi:10.1053/j.ajkd.2013.11.027

17. Eltzschig HK, Sitkovsky MV, Robson SC (2012) Purinergic signaling during inflammation. N Engl J Med 367 (24):2322–2333. doi:10.1056/NEJMra1205750

18. Bours MJ, Swennen EL, Di Virgilio F, Cronstein BN, Dagnelie PC (2006) Adenosine 5’-triphosphate and adenosine as endogenous signaling molecules in immunity and inflammation. Pharmacol Ther 112 (2):358–404. doi:10.1016/j.pharmthera.2005.04.013

19. Nwafor DC, Chakraborty S, Brichacek AL, Jun S, Gambill CA, Wang W, Engler-Chiurazzi EB, Dakhlallah D, Pinkerton AB, Millan JL, Benkovic SA, Brown CM (2020) Loss of tissue-nonspecific alkaline phosphatase (TNAP) enzyme activity in cerebral microvessels is coupled to persistent neuroinflammation and behavioral deficits in late sepsis. Brain Behav Immun 84:115–131. doi:10.1016/j.bbi.2019.11.016

20. Fonta C, Barone P, Rodriguez Martinez L, Negyessy L (2015) Rediscovering TNAP in the Brain: A Major Role in Regulating the Function and Development of the Cerebral Cortex. Subcell Biochem 76:85–106. doi:10.1007/978-94-017-7197-9_5

21. Fonta C, Negyessy L, Renaud L, Barone P (2005) Postnatal development of alkaline phosphatase activity correlates with the maturation of neurotransmission in the cerebral cortex. J Comp Neurol 486 (2):179–196. doi:10.1002/cne.20524

22. Teriete P, Pinkerton AB, Cosford ND (2013) Inhibitors of tissue-nonspecific alkaline phosphatase (TNAP): from hits to leads. Methods Mol Biol 1053:85–101. doi:10.1007/978-1-62703-562-0_5

23. Pinkerton AB, Sergienko E, Bravo Y, Dahl R, Ma CT, Sun Q, Jackson MR, Cosford NDP, Millan JL (2018) Discovery of 5-((5-chloro-2-methoxyphenyl)sulfonamido)nicotinamide (SBI-425), a potent and orally bioavailable tissue-nonspecific alkaline phosphatase (TNAP) inhibitor. Bioorg Med Chem Lett 28 (1):31–34. doi:10.1016/j.bmcl.2017.11.024

24. Foster BL, Kuss P, Yadav MC, Kolli TN, Narisawa S, Lukashova L, Cory E, Sah RL, Somerman MJ, Millan JL (2017) Conditional Alpl Ablation Phenocopies Dental Defects of Hypophosphatasia. J Dent Res 96 (1):81–91. doi:10.1177/0022034516663633

25. Weksler BB, Subileau EA, Perriere N, Charneau P, Holloway K, Leveque M, Tricoire-Leignel H, Nicotra A, Bourdoulous S, Turowski P, Male DK, Roux F, Greenwood J, Romero IA, Couraud PO (2005) Blood-brain barrier-specific properties of a human adult brain endothelial cell line. FASEB J 19 (13):1872–1874. doi:10.1096/fj.04-3458fje

26. Alva JA, Zovein AC, Monvoisin A, Murphy T, Salazar A, Harvey NL, Carmeliet P, Iruela-Arispe ML (2006) VE-Cadherin-Cre-recombinase transgenic mouse: a tool for lineage analysis and gene deletion in endothelial cells. Dev Dyn 235 (3):759–767. doi:10.1002/dvdy.20643

27. Madisen L, Zwingman TA, Sunkin SM, Oh SW, Zariwala HA, Gu H, Ng LL, Palmiter RD, Hawrylycz MJ, Jones AR, Lein ES, Zeng H (2010) A robust and high-throughput Cre reporting and characterization system for the whole mouse brain. Nat Neurosci 13 (1):133–140. doi:10.1038/nn.2467

28. Brichacek AL, Benkovic SA, Chakraborty S, Nwafor DC, Wang W, Jun S, Dakhlallah D, Geldenhuys WJ, Pinkerton AB, Millan JL, Brown CM (2019) Systemic inhibition of tissue-nonspecific alkaline phosphatase alters the brain-immune axis in experimental sepsis. Sci Rep 9 (1):18788. doi:10.1038/s41598-019-55154-2

29. Doll DN, Engler-Chiurazzi EB, Lewis SE, Hu H, Kerr AE, Ren X, Simpkins JW (2015) Lipopolysaccharide exacerbates infarct size and results in worsened post-stroke behavioral outcomes. Behav Brain Funct 11 (1):32. doi:10.1186/s12993-015-0077-5

30. Ma HW, Ye W, Chen HS, Nie TJ, Cheng LF, Zhang L, Han PJ, Wu XA, Xu ZK, Lei YF, Zhang FL (2017) In-Cell Western Assays to Evaluate Hantaan Virus Replication as a Novel Approach to Screen Antiviral Molecules and Detect Neutralizing Antibody Titers. Front Cell Infect Microbiol 7:269. doi:10.3389/fcimb.2017.00269

31. Engelhardt B, Liebner S (2014) Novel insights into the development and maintenance of the blood-brain barrier. Cell and tissue research 355 (3):687–699. doi:10.1007/s00441-014-1811-2

32. Dahl R, Sergienko EA, Su Y, Mostofi YS, Yang L, Simao AM, Narisawa S, Brown B, Mangravita-Novo A, Vicchiarelli M, Smith LH, O’Neill WC, Millan JL, Cosford ND (2009) Discovery and validation of a series of aryl sulfonamides as selective inhibitors of tissue-nonspecific alkaline phosphatase (TNAP). J Med Chem 52 (21):6919–6925. doi:10.1021/jm900383s

33. Deracinois B, Lenfant AM, Dehouck MP, Flahaut C (2015) Tissue Non-specific Alkaline Phosphatase (TNAP) in Vessels of the Brain. Subcell Biochem 76:125–151. doi:10.1007/978-94-017-7197-9_7

34. Debray J, Chang L, Marques S, Pellet-Rostaing S, Le Duy D, Mebarek S, Buchet R, Magne D, Popowycz F, Lemaire M (2013) Inhibitors of tissue-nonspecific alkaline phosphatase: design, synthesis, kinetics, biomineralization and cellular tests. Bioorg Med Chem 21 (24):7981–7987. doi:10.1016/j.bmc.2013.09.053

35. Hall A (1998) Rho GTPases and the actin cytoskeleton. Science 279 (5350):509–514. doi:10.1126/science.279.5350.509

36. Xing L, Remick DG (2005) Mechanisms of dimethyl sulfoxide augmentation of IL-1 beta production. J Immunol 174 (10):6195–6202. doi:10.4049/jimmunol.174.10.6195

37. Koch JC, Tatenhorst L, Roser AE, Saal KA, Tonges L, Lingor P (2018) ROCK inhibition in models of neurodegeneration and its potential for clinical translation. Pharmacol Ther 189:1–21. doi:10.1016/j.pharmthera.2018.03.008

38. M.J. Weiss KR, P.S. Henthorn, B. Lamb, T. Kadesch, H. Harris (1988) Structure of the human liver/bone/kidney alkaline phosphatase gene. JOURNAL OF BIOLOGICAL CHEMISTRY 263:12002–12010

39. Yang J, Ruan F, Zheng Z (2018) Ripasudil Attenuates Lipopolysaccharide (LPS)-Mediated Apoptosis and Inflammation in Pulmonary Microvascular Endothelial Cells via ROCK2/eNOS Signaling. Med Sci Monit 24:3212–3219. doi:10.12659/MSM.910184

40. Grothaus JS, Ares G, Yuan C, Wood DR, Hunter CJ (2018) Rho kinase inhibition maintains intestinal and vascular barrier function by upregulation of occludin in experimental necrotizing enterocolitis. Am J Physiol Gastrointest Liver Physiol 315 (4):G514–G528. doi:10.1152/ajpgi.00357.2017

41. Feng S, Zou L, Wang H, He R, Liu K, Zhu H (2018) RhoA/ROCK-2 Pathway Inhibition and Tight Junction Protein Upregulation by Catalpol Suppresses Lipopolysaccaride-Induced Disruption of Blood-Brain Barrier Permeability. Molecules 23 (9). doi:10.3390/molecules23092371

42. Barone FC, Arvin B, White RF, Miller A, Webb CL, Willette RN, Lysko PG, Feuerstein GZ (1997) Tumor necrosis factor-alpha. A mediator of focal ischemic brain injury. Stroke 28 (6):1233–1244. doi:10.1161/01.str.28.6.1233

43. Michie HR, Manogue KR, Spriggs DR, Revhaug A, O’Dwyer S, Dinarello CA, Cerami A, Wolff SM, Wilmore DW (1988) Detection of circulating tumor necrosis factor after endotoxin administration. N Engl J Med 318 (23):1481–1486. doi:10.1056/NEJM198806093182301

44. Romero CR, Herzig DS, Etogo A, Nunez J, Mahmoudizad R, Fang G, Murphey ED, Toliver-Kinsky T, Sherwood ER (2010) The role of interferon-gamma in the pathogenesis of acute intra-abdominal sepsis. J Leukoc Biol 88 (4):725–735. doi:10.1189/jlb.0509307

45. Yilmaz G, Arumugam TV, Stokes KY, Granger DN (2006) Role of T lymphocytes and interferon-gamma in ischemic stroke. Circulation 113 (17):2105–2112. doi:10.1161/CIRCULATIONAHA.105.593046

46. Mehra A, Guerit S, Macrez R, Gosselet F, Sevin E, Lebas H, Maubert E, De Vries HE, Bardou I, Vivien D, Docagne F (2020) Nonionotropic Action of Endothelial NMDA Receptors on Blood-Brain Barrier Permeability via Rho/ROCK-Mediated Phosphorylation of Myosin. J Neurosci 40 (8):1778–1787. doi:10.1523/JNEUROSCI.0969-19.2019

47. Schubert-Unkmeir A, Konrad C, Slanina H, Czapek F, Hebling S, Frosch M (2010) Neisseria meningitidis induces brain microvascular endothelial cell detachment from the matrix and cleavage of occludin: a role for MMP-8. PLoS Pathog 6 (4):e1000874. doi:10.1371/journal.ppat.1000874

48. Greene C, Hanley N, Campbell M (2019) Claudin-5: gatekeeper of neurological function. Fluids Barriers CNS 16 (1):3. doi:10.1186/s12987-019-0123-z

49. Esue O, Carson AA, Tseng Y, Wirtz D (2006) A direct interaction between actin and vimentin filaments mediated by the tail domain of vimentin. J Biol Chem 281 (41):30393–30399. doi:10.1074/jbc.M605452200

50. Svitkina TM, Verkhovsky AB, Borisy GG (1996) Plectin sidearms mediate interaction of intermediate filaments with microtubules and other components of the cytoskeleton. J Cell Biol 135 (4):991–1007. doi:10.1083/jcb.135.4.991

51. Hollenbeck PJ, Bershadsky AD, Pletjushkina OY, Tint IS, Vasiliev JM (1989) Intermediate filament collapse is an ATP-dependent and actin-dependent process. J Cell Sci 92 (Pt 4):621–631

52. Dupin I, Sakamoto Y, Etienne-Manneville S (2011) Cytoplasmic intermediate filaments mediate actin-driven positioning of the nucleus. J Cell Sci 124 (Pt 6):865–872. doi:10.1242/jcs.076356

53. Jiu Y, Lehtimaki J, Tojkander S, Cheng F, Jaalinoja H, Liu X, Varjosalo M, Eriksson JE, Lappalainen P (2015) Bidirectional Interplay between Vimentin Intermediate Filaments and Contractile Actin Stress Fibers. Cell Rep 11 (10):1511–1518. doi:10.1016/j.celrep.2015.05.008

54. Eriksson JE, Dechat T, Grin B, Helfand B, Mendez M, Pallari HM, Goldman RD (2009) Introducing intermediate filaments: from discovery to disease. J Clin Invest 119 (7):1763–1771. doi:10.1172/JCI38339

55. Yang L, Tang L, Dai F, Meng G, Yin R, Xu X, Yao W (2017) Raf-1/CK2 and RhoA/ROCK signaling promote TNF-alpha-mediated endothelial apoptosis via regulating vimentin cytoskeleton. Toxicology 389:74–84. doi:10.1016/j.tox.2017.07.010

56. Trindade P, Loiola EC, Gasparotto J, Ribeiro CT, Cardozo PL, Devalle S, Salerno JA, Ornelas IM, Ledur PF, Ribeiro FM, Ventura ALM, Moreira JCF, Gelain DP, Porciuncula LO, Rehen SK (2020) Short and long TNF-alpha exposure recapitulates canonical astrogliosis events in human-induced pluripotent stem cells-derived astrocytes. Glia 68 (7):1396–1409. doi:10.1002/glia.23786

57. Shi J, Wu X, Surma M, Vemula S, Zhang L, Yang Y, Kapur R, Wei L (2013) Distinct roles for ROCK1 and ROCK2 in the regulation of cell detachment. Cell Death Dis 4:e483. doi:10.1038/cddis.2013.10

58. Shibuya M, Hirai S, Seto M, Satoh S, Ohtomo E, Fasudil Ischemic Stroke Study G (2005) Effects of fasudil in acute ischemic stroke: results of a prospective placebo-controlled double-blind trial. J Neurol Sci 238 (1-2):31–39. doi:10.1016/j.jns.2005.06.003

59. Fukuta T, Asai T, Yanagida Y, Namba M, Koide H, Shimizu K, Oku N (2017) Combination therapy with liposomal neuroprotectants and tissue plasminogen activator for treatment of ischemic stroke. FASEB J 31 (5):1879–1890. doi:10.1096/fj.201601209R

60. Liu K, Li Z, Wu T, Ding S (2011) Role of rho kinase in microvascular damage following cerebral ischemia reperfusion in rats. Int J Mol Sci 12 (2):1222–1231. doi:10.3390/ijms12021222

61. Zhu J, Zhou B, Ma L, Liu L (2021) Exploring the beneficial role of ROCK inhibitors in sepsis-induced cerebral and cognitive injury in rats. Fundam Clin Pharmacol. doi:10.1111/fcp.12645

