## Supplementary figures for "Brain endothelial cell tissue-nonspecific alkaline phosphatase (TNAP) activity promotes maintenance of barrier integrity *via* the ROCK pathway"

Supplementary Figure 1

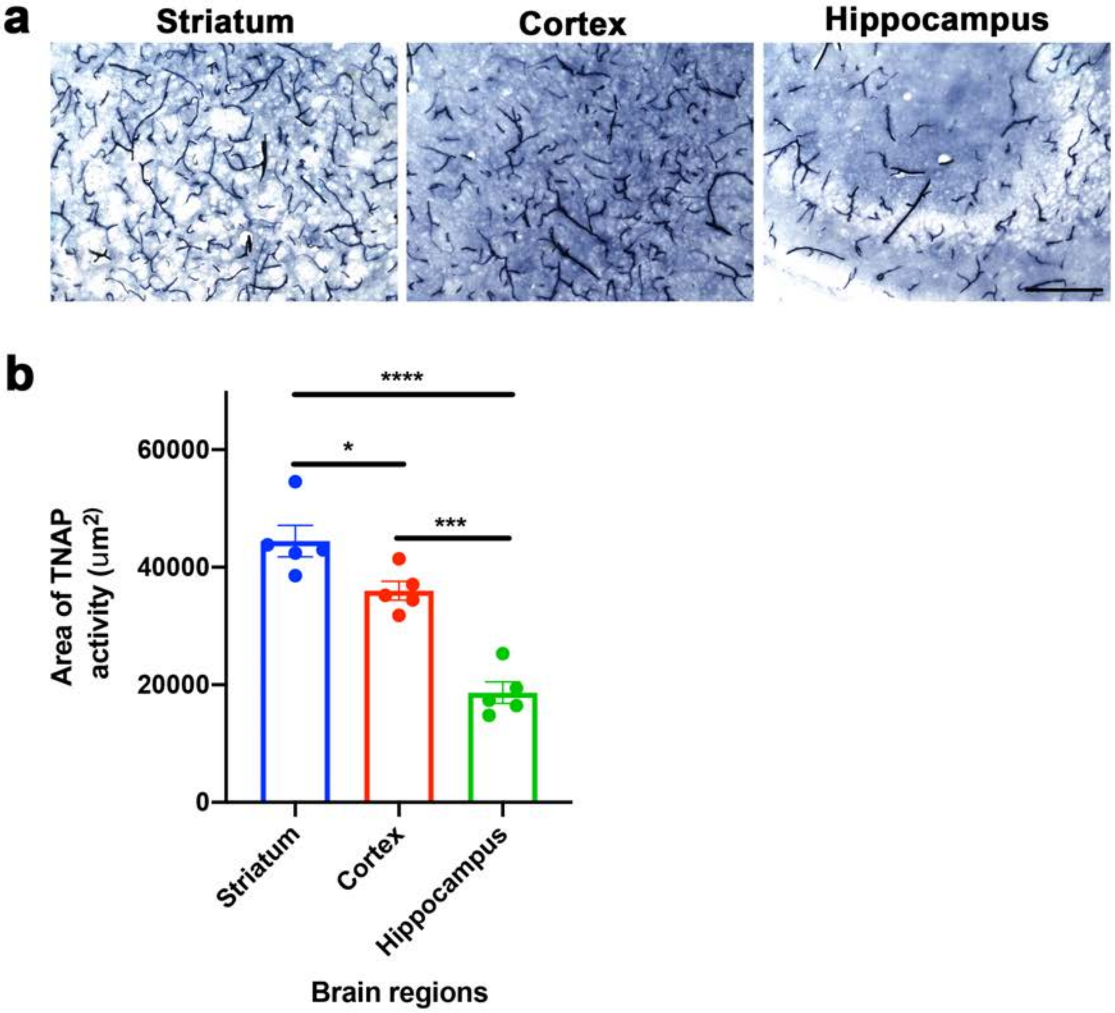

Supplementary Figure 2

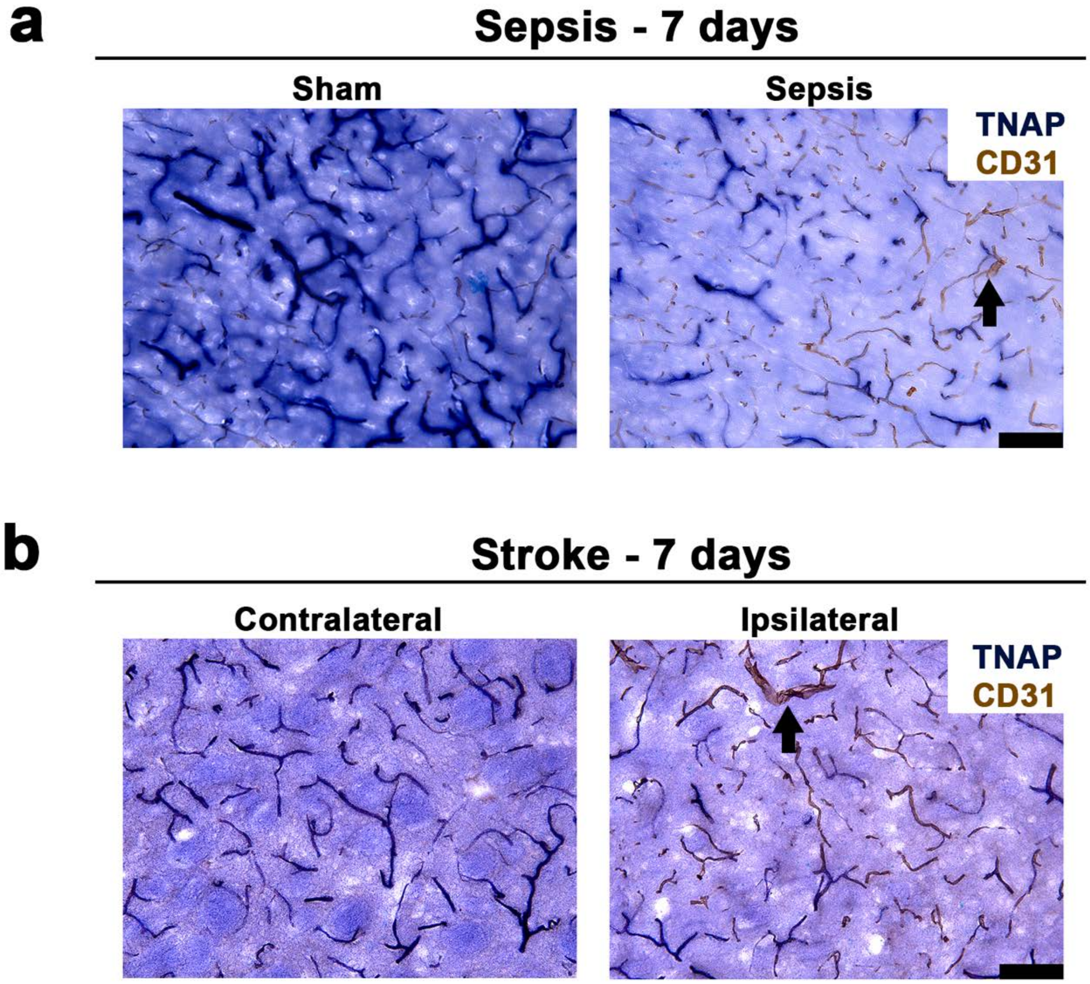

Supplementary Figure 3

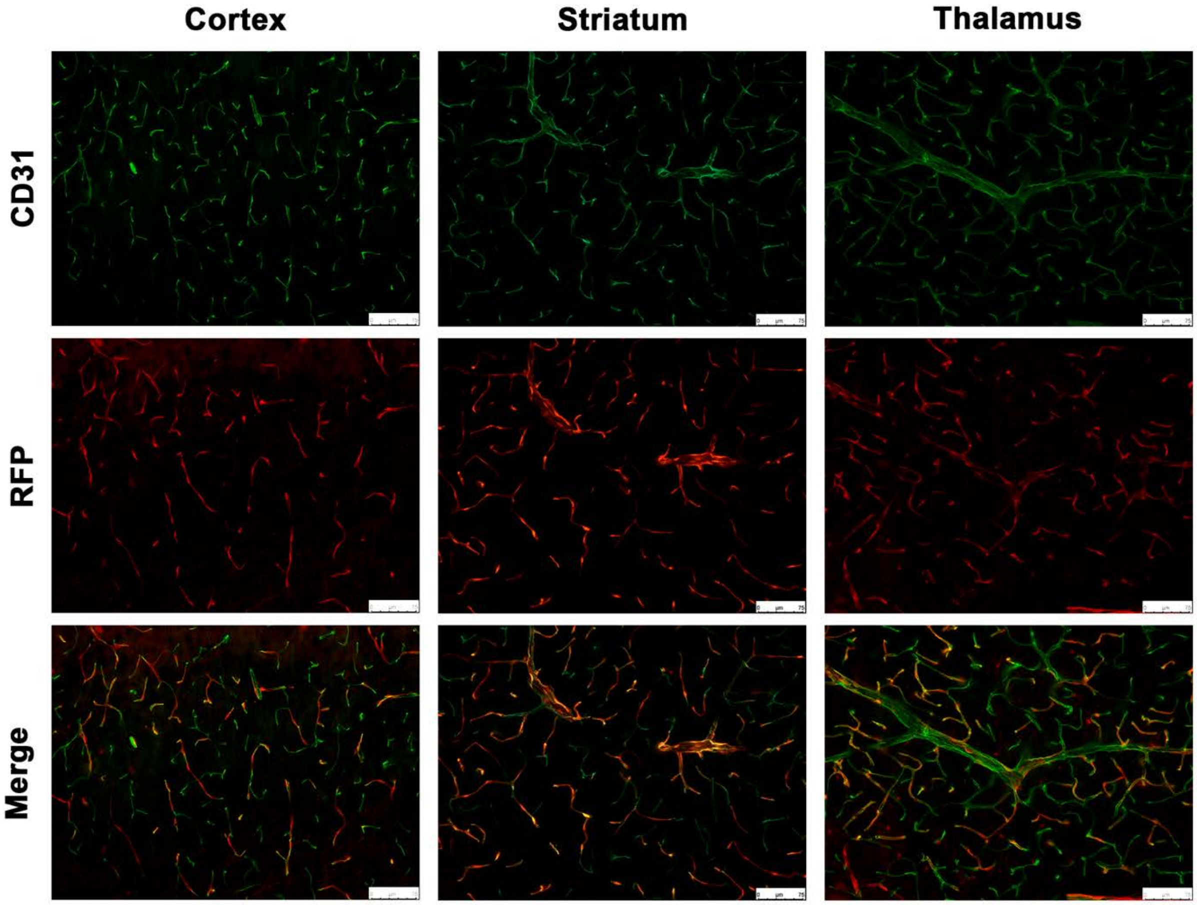

**Supplementary Figure Legends**

**Supplementary Fig. 1** Expression of brain microvascular TNAP activity is region specific in mice. (**a**) Representative images of brain microvascular TNAP activity expression in the cortex, striatum, and hippocampus (CA3 shown) of WT mice (n = 5). (**b**) Quantification of brain microvascular TNAP activity expression revealed that TNAP activity is most significantly expressed in the striatum ($p < 0.0001$, Tukey's multiple comparisons test, one-way ANOVA) and cortex ($p = 0.0002$, Tukey's multiple comparisons test, one-way ANOVA) relative to the hippocampus. Interestingly, TNAP activity in the striatum was significantly increased ($p = 0.03$) when compared to the cortex. * indicates $p < 0.05$, ***$p < 0.001$, and ****$p < 0.0001$, and is considered significant. All data are presented as mean ± SEM. Images taken at 20X magnification and scale bar = 200 μm

**Supplementary Fig. 2** Loss of TNAP activity on brain microvessels. (**a**) Representative images showed loss of TNAP activity on CD31 positive (black arrow) brain microvessels 7 days post-sepsis. (**b**) Similarly, TNAP activity on CD31 positive brain microvessels decreased in the ipsilateral striatal penumbra (stroke hemisphere) compared to the contralateral striatum (non-stroke hemisphere) 7 days post-stroke. Images taken at 20X magnification and scale bar = 250 μm

**Supplementary Fig. 3** Ve-Cadherin Cre is specific for microvessels. Red fluorescent protein (RFP) signal colocalizes with CD31 vascular endothelial marker. Cortex, striatum, and thalamus are shown. Images taken at 20X magnification and scale bar = 75 μm
